# Structural disruption of BAF chromatin remodeller impairs neuroblastoma metastasis by reverting an invasiveness epigenomic program

**DOI:** 10.1101/2022.05.17.492122

**Authors:** Carlos Jiménez, Roberta Antonelli, Mariona Nadal-Ribelles, Laura Devis-Jauregui, Pablo Latorre, Carme Solé, Marc Masanas, Adrià Molero-Valenzuela, Aroa Soriano, Josep Sánchez de Toledo, David Llobet-Navas, Josep Roma, Francesc Posas, Eulàlia de Nadal, Soledad Gallego, Lucas Moreno, Miguel F. Segura

**Affiliations:** Group of Childhood Cancer and Blood Disorders, Vall d’Hebron Research Institute (VHIR), Universitat Autònoma de Barcelona (UAB), Barcelona, Spain; Institute for Research in Biomedicine, The Barcelona Institute of Science and Technology, Barcelona, Spain; Department of Medicine and Life Sciences (MELIS), Universitat Pompeu Fabra (UPF), Barcelona, Spain; Molecular Mechanisms and Experimental Therapy in Oncology-Oncobell Program, Bellvitge Biomedical Research Institute, L’Hospitalet de Llobregat, Spain; Catalan Institute of Oncology, L’Hospitalet de Llobregat, Spain; Low Prevalence Tumors. Centro de Investigación Biomédica en Red de Cáncer (CIBERONC), Instituto de Salud Carlos III, Madrid, Spain; Paediatric Oncology and Haematology Department, Vall d’Hebron University Hospital, Barcelona, Spain

**Keywords:** Epigenetics, Epigenomics, Cancer, Neuroblastoma, Chromatin remodelling, SWI/SNF, Metastasis

## Abstract

**Background:** Epigenetic programming during development is essential for determining cell lineages, and alterations in this programming contribute to the initiation of embryonal tumour development. In neuroblastoma, neural crest progenitors block their course of natural differentiation into sympathoadrenergic cells, leading to the development of aggressive and metastatic paediatric cancer. Research of the epigenetic regulators responsible for oncogenic epigenomic networks is crucial for developing new epigenetic-based therapies against these tumours. Mammalian switch/sucrose non-fermenting (mSWI/SNF) ATP-dependent chromatin remodelling complexes act genome-wide translating epigenetic signals into open chromatin states. The present study aimed to understand the contribution of mSWI/SNF to the oncogenic epigenomes of neuroblastoma and its potential as a therapeutic target.

**Methods:** Functional characterisation of the mSWI/SNF complexes was performed in neuroblastoma cells using proteomic approaches, loss-of-function experiments, transcriptome and chromatin accessibility analyses, and *in vitro* and *in vivo* assays.

**Results:** Neuroblastoma cells contain three main mSWI/SNF subtypes, but only BRG1-associated factor (BAF) complex disruption through silencing of its key structural subunits, ARID1A and ARID1B, impairs cell proliferation by promoting cell cycle blockade. Genome-wide chromatin remodelling and transcriptomic analyses revealed that BAF disruption results in the epigenetic repression of an extensive invasiveness-related expression program involving integrins, cadherins, and key mesenchymal regulators, thereby reducing adhesion to the extracellular matrix and the subsequent invasion *in vitro* and drastically inhibiting the initiation and growth of neuroblastoma metastasis *in vivo*.

**Conclusions:** We report a novel ATPase-independent role for the BAF complex in maintaining an epigenomic program that allows neuroblastoma invasiveness and metastasis, urging for the development of new BAF pharmacological structural disruptors for therapeutic exploitation in metastatic neuroblastoma.

## Background

Epigenetic control of gene expression is essential for the determination and maintenance of the different developmental cell lineages that must coexist in the tissues of a newly forming organism. Epigenetic alterations during development contribute to the development of embryonal tumours, originating from errors in the regular developmental process that lead to aberrant undifferentiated and proliferating cell populations [1]. Neuroblastoma, an embryonal tumour of the sympathetic nervous system, originates from undifferentiated precursor cells of the neural crest aimed at undergoing dorsolateral migration and becoming sympathoadrenergic cells. These cells fail to undergo a complete adrenergic differentiation course and initiate a neoplastic process, leading to the onset of an aggressive and metastatic paediatric oncologic disease [2]. Molecular alterations driving neuroblastoma initiation and progression include aberrant epigenetic landscapes, such as abnormal DNA methylation patterns [3,4] or oncogenic super-enhancers that determine malignant phenotypes [5,6].

Chromatin remodelling is the final step in epigenetic regulation through which specific chromatin condensation states are achieved. It is the final consequence of the cascade of molecular events triggered by specific epigenetic signals (i.e. DNA methylation, histone marks, or chromatin-associated non-coding RNA), followed by their recognition by epigenetic readers, and culminating in the activity of a chromatin remodeller at a specific genomic locus [7]. ATP-dependent chromatin remodellers are molecular machines that use ATP hydrolysis as the energy source to produce diverse effects on the chromatin structure. Among them, mammalian switch/sucrose non-fermenting (mSWI/SNF) remodellers are huge macromolecular complexes that increase chromatin accessibility by generating nucleosome-depleted regions [8], thereby enabling the direct interaction between DNA and certain regulatory proteins, usually transcription factors [9,10]. These remodellers act at a genome-wide level [11,12] and have tissue-specific regulatory functions [13,14], which makes them relevant epigenetic coactivators that translate epigenetic signals into open chromatin states and help maintain cell identity. mSWI/SNF are multiprotein macro-complexes that exist in three different intracellular coexistent subtypes, namely, *BRG1-Associated Factor* (BAF), *Polybromo-associated BAF* (PBAF), and *non-canonical BAF* (ncBAF) complexes, all of which have specific subunit compositions and divergent genomic occupancies [15–17].

The special relevance of the mSWI/SNF complexes in cancer, reported in the early 2000s and in continued increase in the last two decades, is reflected in the fact that ~20% of human cancers harbour deleterious mutations in at least one mSWI/SNF subunit, a mutation rate comparable to that of well-known driving tumour suppressors such as *TP53* and *PTEN* [18]. Moreover, aberrations involving certain mSWI/SNF subunits are defining and ubiquitous traits of certain rare tumours of embryonal origin, such as *SMARCB1* loss in rhabdoid tumours [19], or translocations involving the *SS18* gene in synovial sarcoma [20]. However, oncogenic functions of this complex have been reported in different human malignancies, with certain oncogenic functions being dependent on the expression of determined mSWI/SNF subunits, and their overexpression correlating with poor prognosis in patients [21–24].

The embryonal origin of neuroblastoma, together with the driving role of epigenetic alterations in these tumours, put mSWI/SNF complexes in the spotlight as plausible executors of the aberrant epigenetic landscapes by translating them into chromatin states and maintaining specific oncogenic transcriptional programs. Indeed, deleterious mutations in the genes encoding the mSWI/SNF subunits ARID1A and ARID1B have been reported in neuroblastoma samples and are associated with poor patient prognosis [25,26]. Moreover, ARID1A loss has functional driving effects on the oncogenic functions of neuroblastoma cells, including invasiveness and adrenergic-to-mesenchymal transition [27,28]. Conversely, the mRNA and protein expression of one of the catalytic subunits, BRG1 (or SMARCA4), was correlated with advanced stages and poor prognosis in neuroblastoma patients, and this subunit was reported to control an oncogenic transcriptional program in neuroblastoma cells [29].

Hence, an in-depth study of this chromatin remodeller as a whole, and not of its subunits as independent entities, is still needed to elucidate its complete set of functions in neuroblastoma. We present a systematic analysis of the presence, composition, and role of mSWI/SNF in neuroblastoma cells, which led to the discovery of multiple cancer-relevant transcriptional regulatory functions attributed exclusively to the structural integrity of the BAF-subtype complex. Structural disruption of the BAF complex, achieved by silencing of its essential subunits ARID1A and ARID1B, exerted an epigenomic reprogramming in neuroblastoma cells, resulting in the repression of a wide expression program involving invasiveness-related genes such as integrins, cadherins, and key mesenchymal regulators, leading to the impairment of metastatic invasion, an effect that is potentially exploitable in a therapeutic strategy for the treatment of metastatic neuroblastoma.

## Methods

### Proliferation, adhesion, and invasion assays

Neuroblastoma proliferation experiments were performed by seeding 8 × 10^4^ cells/well in 24-well plates and allowing them to grow for 72 h. The cells were fixed with 1% glutaraldehyde and stained with 0.5% crystal violet. Dried crystals were dissolved in 15% acetic acid and the absorbance was measured at 590 nm. For adhesion assays, 96-well culture plates were pre-coated with poly-D-lysine (Sigma-Aldrich, St. Louis, MO, USA; 0.5 mg/mL in water, 20 min at 37 °C), followed by coating with collagen I (Corning Inc., Corning, NY, USA; 80 μg/mL in 0.02 M acetic acid, overnight evaporation), and 4 × 10^4^ cells/well were seeded. Non-adherent cells were removed in a time course of 5-minute intervals by aspiration. The remaining collagen-adhered cells were allowed to stand for 8 h in full medium for complete attachment; afterwards, the cells were fixed with 1% glutaraldehyde and stained with crystal violet. Adhesion percentage was normalised to empty wells (0% adhesion) and non-aspirated cells (100% adhesion). Invasion assays were performed by seeding 2 × 10^5^ cells in serum-free media in the upper chamber of transwells (Corning; 8.0 μm pore size) previously coated with a collagen barrier, upon a lower chamber filled with serum-containing media. After 16 h, the remaining cells were removed from the upper chamber, and the cells that had migrated to the lower membrane surface were fixed with 4%paraformaldehyde and stained with crystal violet. Invading cells were imaged by bright field microscopy, quantified by diluting crystals in acetic acid, and the absorbance was read at 590 nm.

### RNA-sequencing

RNA was purified from neuroblastoma cells using Qiazol buffer and the miRNeasy Mini Kit (Qiagen, Hilden, Germany), following the manufacturer’s instructions. Total RNA was eluted in nuclease-free water and fluorescently quantified using the Qubit RNA HS Assay (Invitrogen, Waltham, MA, USA), and its quality was evaluated with the RNA 6000 Nano Assay on a Bioanalyzer 2100 (Agilent, Santa Clara, CA, USA). Libraries were prepared using the TruSeq Stranded mRNA LT Sample Prep Kit protocol (Illumina, San Diego, CA, USA) and sequenced on NovaSeq 6000 (Illumina) at the National Center of Genomic Analyses (CNAG-CRG, Barcelona, Spain). For detailed procedures and analysis, see Supplementary Materials and Methods (Additional file 1).

### ATAC-sequencing

ATAC-seq samples were processed in triplicate as previously reported [30], with minor modifications. Transposase Tn5 E54K, L372P was expressed and purified from the pETM11 vector, loaded with linker oligonucleotides as previously reported [31], purified with a 30-kDa Amicon Ultra 0.5 centrifugal unit (Sigma-Aldrich) and diluted to 0.1 mg/mL in 25% glycerol. Nuclei corresponding to 5 × 10^5^ cells were mixed with 5 μL of loaded Tn5 and tagmentation mix, and the mixture was incubated for 60 min at 37 °C. Tn5 was heat-inactivated for 5 min at 80 °C. DNA was purified using the MinElute PCR Purification kit (Qiagen) and PCR-amplified using NEBNext 2× Ultra Mastermix (New England Biolab, Ipswich, MA, USA) with the combinatorial dual index primers i5 and i7 (IDT, Newark, NJ, USA) for 12 cycles. Libraries were purified with Ampure Beads (Beckman Coulter, Brea, CA, USA) and quality-checked with Agilent High-Sensitivity DNA on an Agilent 2100 instrument (Agilent). Samples were sequenced on a NovaSeq 6000 (Illumina) at CNAG-CRG. For detailed procedures and analysis, see Supplementary Materials and Methods (Additional file 1).

### Neuroblastoma metastasis murine models

Experimental procedures involving animals were performed at the Rodent Platform of the Laboratory Animal Service of the Vall d’Hebron Institute of Research (LAS, VHIR, Barcelona, Spain). All animal protocols were reviewed and approved by the regional Institutional Animal Care and Ethics Committee of Animal Experimentation. Murine neuroblastoma metastatic models were generated as previously described [32]. SK-N-BE(2) neuroblastoma cells stably transduced with the lentiviral FLUC vector, originally named CSCW2-Fluc-ImC [33], containing mCherry and Firefly Luciferase reporters in a Fluc-IRES-mCherry expression cassette, and prepared in 200 μL of phosphate-buffered saline (PBS), were intravenously injected into 5-week-old Fox Chase SCID beige mice (Charles River, Wilmington, MA, USA).

For short-term metastasis experiments, 2 million SK-N-BE(2)-FLUC cells were stained immediately before injection with CellTrace™ Far Red (Invitrogen) following the manufacturer’s instructions. At days 4 and 7 after injection, mice were euthanised, following which livers were collected, mechanically dissociated, and incubated with 10 mg/mL collagenase type I (Gibco, Amarillo, TX, USA) under agitation at 37 °C for 30 min. Homogenates were filtered through a 100-μm cell strainer (Corning) and washed with PBS. Red blood cells were eliminated by incubation for 10 min on ice with 5 mL of erythrocyte lysis buffer (Qiagen). Human cells were enriched using the Mouse Cell Depletion Kit and the QuadroMACS magnetic cell separator (Miltenyi Biotec, Bergisch Gladbach, Germany). Pairs of liver samples from the same experimental group were pooled and subjected to a second round of purification. Eluates were stained with SYTOX Blue (Invitrogen) and analysed using the fluorescence-activated cell sorter LSR Fortessa (BD Biosciences, Franklin Lakes, NJ, USA) at the Flow Cytometry facility of VHIR. The results were analysed using FlowJo v10.8 software (BD Biosciences).

For long-term metastasis experiments, 2 × 10^5^ SK-N-BE(2)-FLUC cells were injected into the tail vein. At the indicated time points, the mice were anaesthetised by isoflurane inhalation and intraperitoneally injected with D-luciferine (Invitrogen). Luminescence signals were acquired using an IVIS SpectrumCT In Vivo Imaging System (PerkinElmer, Waltham, MA, USA), and visualised and quantified using Living Image Software 4.5.2 (Perkin Elmer). Average photon counts from same-duration expositions were calculated for the whole body for each mouse and normalised by extracting the background signal. Mice were monitored for symptomatology such as signs of suffering or appearance of palpable metastatic lesions, as endpoint criteria.

## Results

### Proliferation of neuroblastoma cells depends on BAF complex structural integrity

We initiated a systematic study on mSWI/SNF in neuroblastoma by determining the structural integrity and composition of its variants. Isolation of all complex subtypes existing in neuroblastoma cell lines by immunoprecipitation of the ubiquitous BRG1 catalytic subunit and analysis of its interactors by mass spectrometry led to the identification of all known mSWI/SNF subunits from the core and ATPase modules, as well as those specific to the three same-cell coexistent variants (Figure 1A, Table S1), except from BRM (or SMARCA2), which is mutually exclusive with BRG1, the bait of the proteomic study. This reflects the presence and structural integrity of the three known variants in neuroblastoma cells. To elucidate the contribution of the mSWI/SNF complex to the oncogenic properties of neuroblastoma cells, we tested the effects of different loss-of-function strategies.

**Fig. 1:**
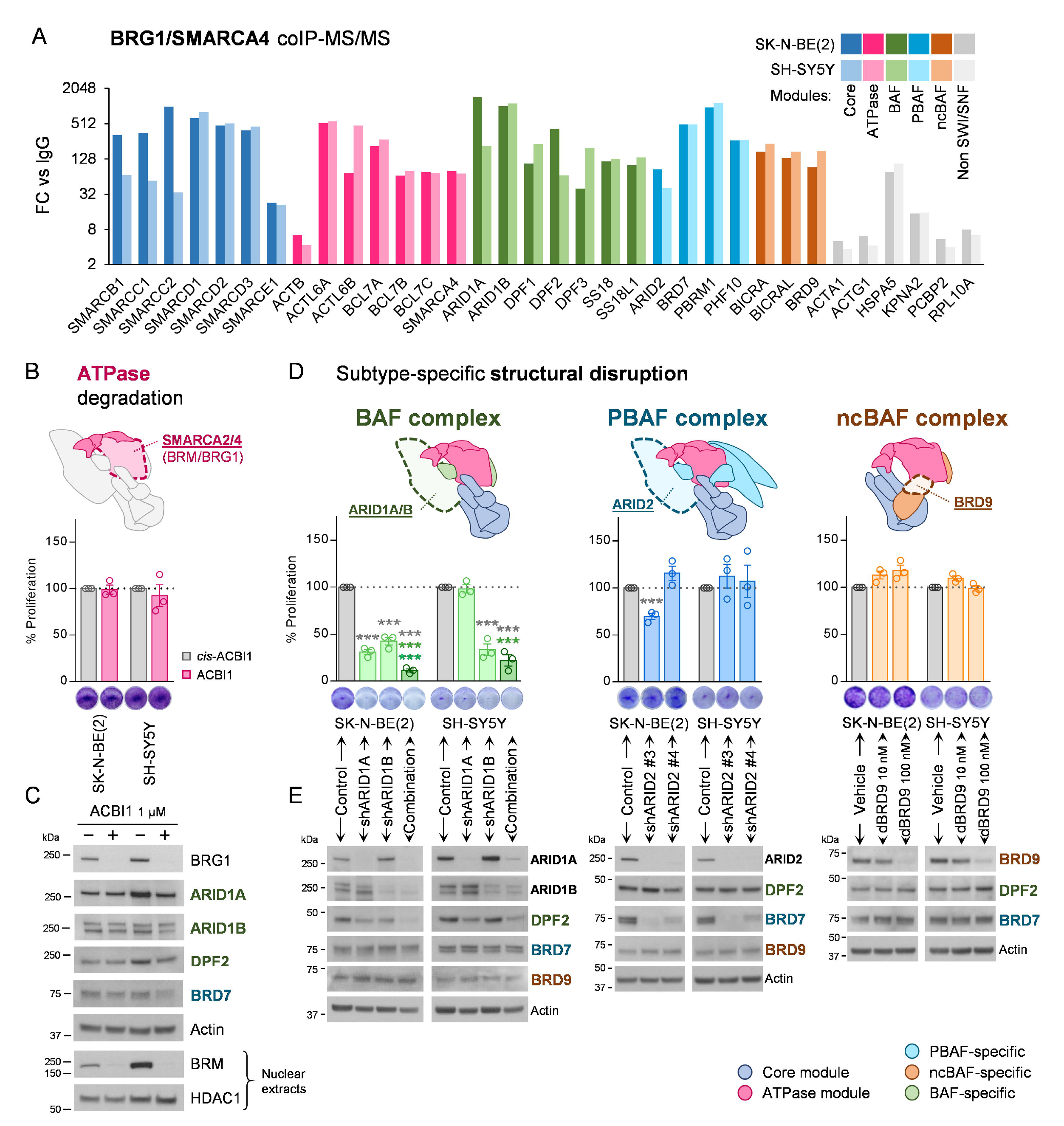
Structural integrity of BAF complex is essential for neuroblastoma proliferation. (**A**) Mass spectrometry analysis of mSWI/SNF complexes composition in SK-N-BE(2) and SH-SY5Y neuroblastoma cell lines. Proteins enriched in BRG1/SMARCA4 co-immunoprecipitation compared with normal IgG control in both cell lines (Bayesian False Discovery Rate (BDFR) < 0.05) are shown. Fold change of average spectrum counts with respect to IgG (FC vs IgG) is represented. (**B**) Degradation of mSWI/SNF ATPase subunits in neuroblastoma cells. Upper panel, schematic representation of the catalytic subunits BRM/SMARCA2 and BRG1/SMARCA4. Lower panel, proliferation assay of neuroblastoma cell lines treated with 1 μM of the BRM/BRG1 degrader ACBI1, or its negative control *cis*-ACBI1 for 96 h. (**C**) Protein expression analysis of mSWI/SNF subunits in neuroblastoma cells treated with 1 μM *cis*-ACBI1 (labelled as −) or ACBI1 (labelled as +) for 96 h. BRM analysis was performed on nuclear extracts, using HDAC1 as nuclear loading control. (**D**) mSWI/SNF subtypes specific disruption in neuroblastoma cells. Upper panel, schematic representation of the key subtype-specific subunits selected for gene silencing. Lower panel, 96 h proliferation assays with neuroblastoma cell lines seeded at 72 h post-transduction with shRNA against the indicated genes or with a non-silencing shRNA control. For BRD9 degradation, cells were treated with the dBRD9 degrader or vehicle at the indicated doses. (**E**) Protein expression analysis of mSWI/SNF subunits in neuroblastoma cells transduced with shRNA or treated with dBRD9 for 96 h. *** means *p* < 0.001. Colour code indicates the condition against which the multiple comparison statistical tests are calculated.

First, we focused on the catalytic subunits BRM and BRG1, the ATPase activity of which is crucial for active chromatin remodelling by all known mSWI/SNF variants. PROTAC degradation of these two subunits did not exert any effect on cell proliferation (Figure 1B) or on the protein levels of other mSWI/SNF subunits, indicating the permanence of the structural integrity of the different complexes after this inhibitory strategy (Figure 1C). Nevertheless, the subtype-specific structural disruption of each of the three same-cell coexistent mSWI/SNF variants by silencing of known subtype-specific and essential structural subunits [16] showed a clear proliferative dependency of neuroblastoma cells on BAF complex structural integrity, which was observed when silencing the pair of redundant and mutually exclusive BAF-specific subunits ARID1A/B (Figure 1D, Figure S1A-B), and also after ARID1B silencing in the ARID1A-defficient COG-N-278 neuroblastoma cell line (Figure S1C-D). This dependency was not reported for the PBAF complex, through ARID2 silencing, or for ncBAF, through BRD9 degradation (Figure 1D). Monitoring each of the subtype-specific subunits, DPF2, BRD7, and BRD9 for BAF, PBAF, and ncBAF, respectively, showed their specific destabilisation after each silencing experiment, thereby confirming the efficacy and specificity of the structural disruption of each variant (Figure 1E). The relevance of the structural integrity of the mSWI/SNF complex on neuroblastoma proliferative function was further proven by the simultaneous full disassembly of the three complex variants by silencing of the SMARCC1 and SMARCC2 core subunits (Figure S1C-D), which are essential for the initiation of the assembly process [16].

These results show that, among the three coexistent and fully assembled mSWI/SNF complexes found in neuroblastoma cells, only the structural integrity of the BAF complex, and not its ATPase activity, is important for neuroblastoma proliferation.

### Structural disruption of the BAF complex by ARID1A and ARID1B silencing exerts an extensive impact on the neuroblastoma transcriptome involving cell cycle blockade

The marked effect exerted by BAF complex disruption in comparison with that of PBAF and ncBAF encouraged us to break down the detailed functions of this variant in neuroblastoma. Because the main functional difference between the complex subtypes is their genomic occupancy and the transcriptional network under their specific control [17,34], we sought to determine the gene expression network under BAF control by performing a transcriptome-wide analysis. RNA-Seq of neuroblastoma cells after BAF disruption by silencing of ARID1A and ARID1B with two different shRNAs for each protein (Figure S2A) revealed the presence of hundreds of transcripts whose expression was modulated after the simultaneous knockdown of both subunits. Contrarily, single silencing of each subunit showed smaller and subunit-specific groups of modulated genes, the majority of which were also modulated after complete disruption of the complex (Figure 2A), illustrating the redundant and compensatory functions of these two subunits in neuroblastoma. Therefore, we focused on the genes that were modulated after the combined silencing of ARID1A and ARID1B, which were named BAF-modulated genes (Figure 2B; Additional file 2). Average expression levels of these genes were more pronouncedly modulated after full BAF disruption than after individual silencing of ARID1A or ARID1B, although these genes were modulated in the same direction by both proteins separately (Figure S2B-C). Of note, sorting this set of BAF-modulated genes according to their behaviour after single inhibitions revealed that the majority of them (50.61%) belonged to clusters of genes regulated by ARID1A and ARID1B separately, but with more pronounced modulation after simultaneous inhibition (Clusters 1 and 5, Figure 2B-C). Comparison between single and combined knockdowns within these clusters revealed that, in all of them, the average expression levels of the included genes was more modulated after full BAF disruption than after single silencing, even in cases in which those genes were modulated by only one of the proteins (Figure 2D). These results showed the synergistic effects of simultaneously inhibiting ARID1A and ARID1B on the transcriptome of these cells in terms of both gene network extension and intensity of gene expression modulation, indicating that only complete disruption of the BAF complex through this combined silencing can uncover all of its transcriptomic functions.

**Fig. 2:**
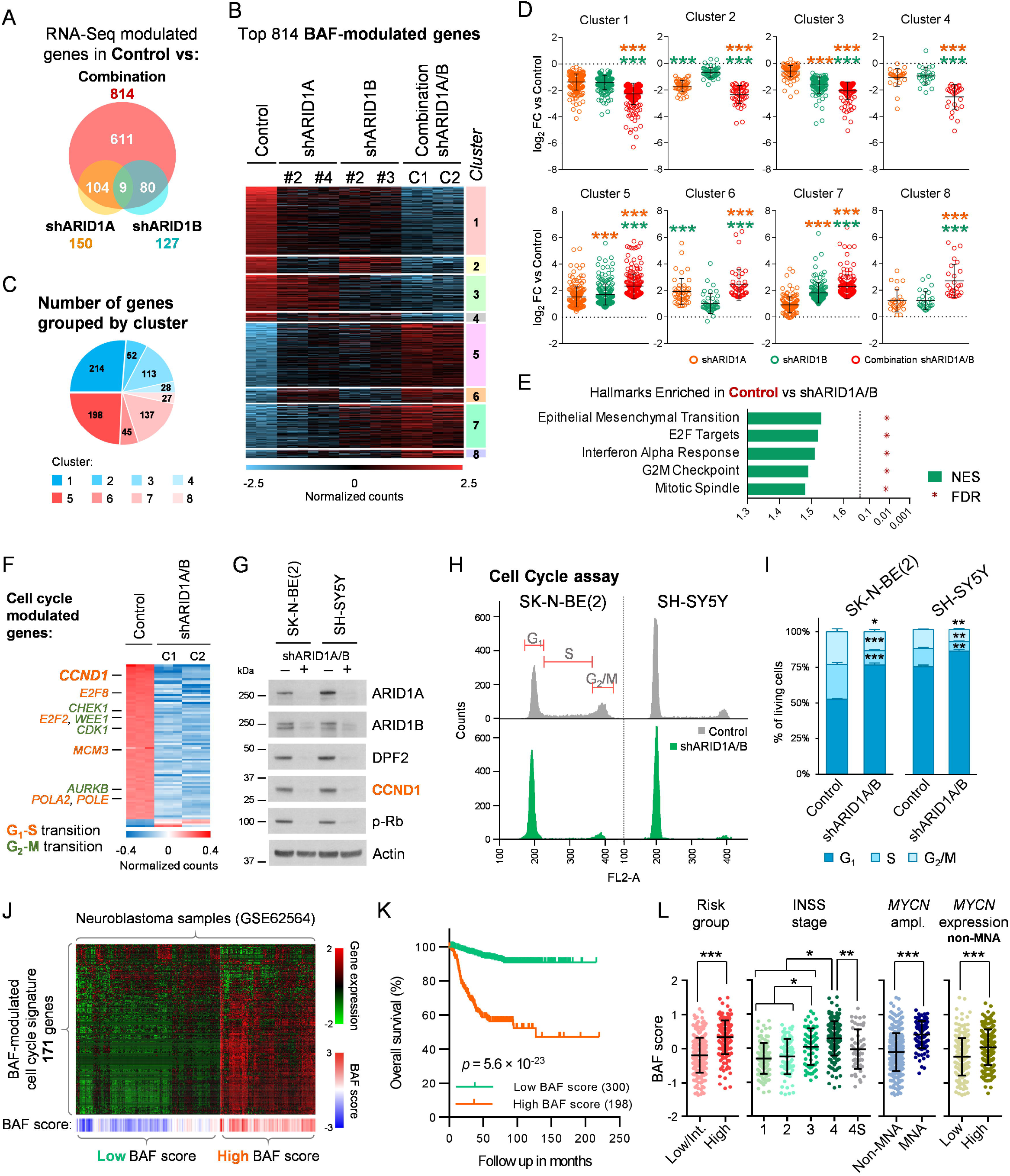
Co-depletion of ARID1A and ARID1B uncovers a wide transcriptome reprogramming involving cell cycle blockade. (**A**) Modulated genes in SK-N-BE(2) cells after single silencing of *ARID1A* or *ARID1B* with two different shRNAs for each gene (shARID1A, shARID1B) and two combinations of shRNA for both genes (Combination) analysed by RNA-Seq. Venn diagram shows the number of modulated genes in comparison with a non-silencing control for each of the three experimental conditions, using cutoff values of log_2_ FC (Fold Change) > 1.5 or < −1.5, and adjusted *p*-value < 0.001. (**B**) Heatmap representing relative RNA expression of genes modulated after BAF-disruption (Combination), using a cutoff of log_2_ FC > 1.5 or < −1.5, and adjusted *p*-value < 0.001. Genes were sorted by single inhibition behaviour-based clusters. (**C**) Pie chart representing the proportion of BAF-modulated genes included in each cluster. (**D**) Comparison of expression FC against control among experimental groups of BAF-modulated genes split in single inhibition behaviour-based clusters. Colour code indicates the condition against which the multiple comparison statistical tests are calculated. (**E)** Normalized Enrichment Score (NES) and False Discovery Rate (FDR) of the top 5 hallmarks of transcriptionally enriched genes in control compared with BAF-depleted cells (shARID1A/B), by Gene Set Enrichment Analysis. (**F**) Heatmap representing relative RNA expression of those genes included in E2F targets, G_2_-M checkpoint and mitotic spindle hallmarks from MSigDB, and modulated after BAF-disruption using a cutoff of log_2_ FC > 1 or < −1, and adjusted *p*-value < 0.001. (**G**) Cyclin D1 (CCND1) and phosphorylated Rb (p-Rb) western blot analysis of BAF-disrupted cells, at 96 h after transduction with shARID1A and shARID1B. (**H**) Cell cycle analysis of neuroblastoma cell lines co-transduced with shARID1A and shARID1B, or with control shRNA, analyzed by flow cytometry. (**I**) Quantification and comparison of the different cell cycle phase populations detected by flow cytometry. (**J**) RNA expression of a BAF-modulated the cell cycle transcriptional signature consisting of 171 genes in a gene expression dataset of 498 human neuroblastoma tumours (GSE62564). Patients were unbiasedly clustered into high and low BAF score groups, as described in Supplementary Materials and Methods. (**K**) Kaplan-Meier plots comparing the overall survival of high and low BAF score group of patients. (**L**) Comparison of BAF signature score according to risk groups, INSS (International Neuroblastoma Staging System) stages, *MYCN* amplification (MNA) and *MYCN* mRNA expression (above or below median) in non-MNA cases. * means *p*< 0.05; ** means *p* < 0.01; *** means *p* < 0.001.

Functional annotation of BAF-modulated genes revealed multiple cell cycle-related gene sets among the most repressed biological hallmarks (Figure 2E, Figure S2D-H, Table S2). These genes, mainly repressed, were implicated both in the progression of the cell cycle from G_1_ to S phase (E2F targets) as well as at later points of the cell cycle (G_2_-M checkpoint, mitotic spindle) (Figure 2F), a repressive event potentially explained by G_1_ phase arrest. Silencing of ARID1A/B caused a strong decrease in cyclin D1 protein levels in neuroblastoma cells, with a concomitant reduction in Rb phosphorylation (Figure 2G). Flow cytometry cell cycle analysis confirmed a clear increase in the percentage of G_1_ phase cells and a reduction in the S and G_2_/M phases, validating cell cycle arrest after BAF complex disruption (Figure 2H-I). No signs of apoptosis or other types of cell death involving cell membrane permeabilisation were observed in the long term (Figure S3), indicating that the decrease in proliferation observed after BAF disruption was related to strong cell cycle arrest, without the contribution of cell death. Interrogation of a cell cycle-related BAF-controlled transcriptional signature, consisting of 171 genes (Table S3), on patient sample expression datasets correlated the BAF activity score with poor prognosis, advanced stages of the disease, *MYCN* amplification and *MYCN* mRNA expression in non-*MYCN* amplified cases (Figure 2J-L, Figure S4). However, this signature retained its survival predictive value when analysing independently *MYCN* non-amplified cases; and after splitting by *MYCN* high and low expression levels in this subset of patients (Figure S4B-D, Table S4). These findings support the idea that increased BAF activity has *MYCN-*related but independent oncogenic consequences in neuroblastoma.

These results prove that BAF complex disruption promotes transcriptome-wide reprogramming of neuroblastoma cells, affecting cell cycle regulators, together with a strong arrest of the G_1_ phase of the cell cycle, thereby blocking neuroblastoma proliferation.

### BAF complex disruption promotes the epigenetic repression of a wide invasiveness-related gene expression network

Given that most of the transcriptome changes occurring after BAF depletion could be attributed to indirect consequences through downstream effectors, we sought to refine the detection of genes under immediate BAF complex epigenetic regulation through *cis* regulatory elements. ATAC-Seq allowed us to identify at a genome-wide level open chromatin regions undergoing significant declines in chromatin accessibility after disruption of this remodeller, the basic function of which is the generation and maintenance of open chromatin states. Among the consensus ATAC-Seq peaks corresponding to the open chromatin sites, more than 19,375 showed significant modulation after BAF disruption, with the majority of them (16,849) being repressive events leading to a close chromatin conformation (Figure 3A, Figure S5A). Peak annotation by genomic proximity identified 7,613 genes associated with these chromatin remodelling repressive events, 719 of which were also transcriptionally modulated in the RNA-Seq data (Figure 3B). Most of these genes (469) underwent significant and consistent transcriptional repression (Figure 3C; Additional file 3). Notably, the cell cycle-related BAF transcriptional signature retained its independent predictive value in neuroblastoma datasets after keeping only those genes associated with chromatin remodelling repressive events (26 genes, Table S5), further associating BAF activity with poor prognosis in patient samples (Figure S6, Table S6). These chromatin remodelling repressive events happening after BAF disruption did not occur at promoters, but rather at both upstream and downstream distal *cis* regulatory elements located in intergenic and intronic regions (Figure 3D, Figure S4B-C). These results are in agreement with the described tendency of the BAF complexes to occupy and regulate enhancers in comparison with PBAF and ncBAF tendencies toward promoters [17,34].

**Fig. 3:**
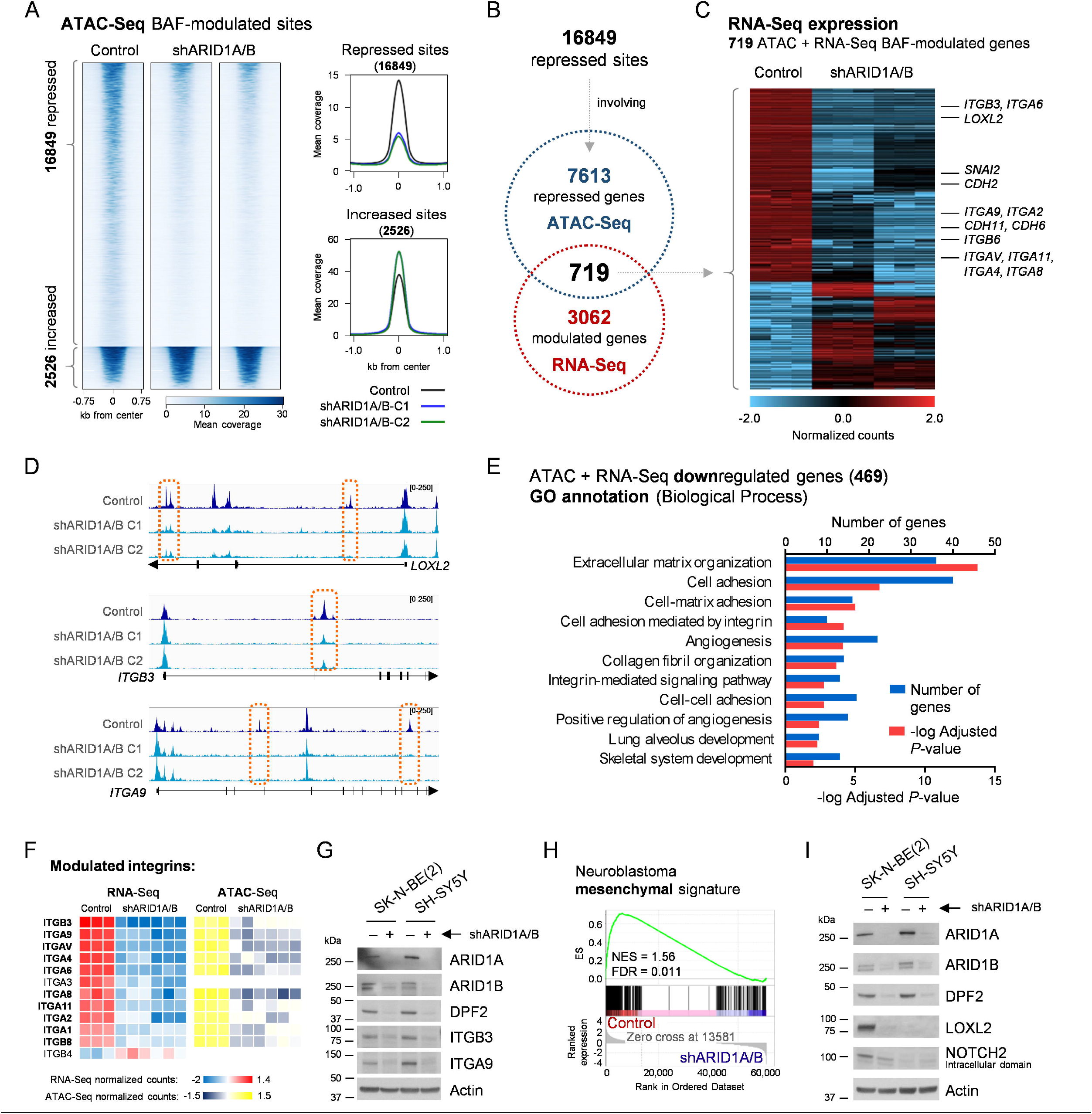
Genome-wide repression of chromatin accessibility after BAF disruption involves an invasiveness-related gene expression network. (**A**) Heatmap and profile plots of significantly ATAC-Seq modulated peaks (False Discovery Rate < 0.01) after BAF disruption (shARID1A/B) in SK-N-BE(2). Two different combinations of shRNAs against both genes were used. (**B**) Comparison of genes annotated to chromatin repressive events by ATAC-Seq with BAF-modulated genes by RNA-Seq (log_2_ FC > 1 or < −1, adjusted *p*-value < 0.05). (**C**) Heatmap representing relative RNA expression of 719 BAF disruption-modulated genes and associated to ATAC-Seq chromatin accessibility repressive events. (**D**) Representative ATAC-Seq coverage images of the genome browser of repressed sites after BAF disruption in the indicated genes also repressed at the mRNA level. (**E**) Gene Ontology (GO) Biological Process analysis of 469 ATAC and RNA-Seq repressed genes, showing enriched categories with adjusted *p*-value < 0.01. (**F**) Heatmap representing relative mRNA expression of 11 BAF-modulated integrin genes and the mean normalized ATAC-Seq counts of their observed chromatin accessibility repressive events. (**G**) Integrin ß3 and α9 western blot analysis in BAF-disrupted cells, at 96 h after co-transduction with shARID1A and shARID1B. (**H**) Gene Set Enrichment Analysis plot of the neuroblastoma mesenchymal phenotype gene expression signature on control versus shARID1A/B (RNA-Seq). Normalized Enrichment Score (NES) and False Discovery Rate (FDR) are shown. (**I**) Western blot validation of the mesenchymal proteins LOXL2 repression after BAF disruption, 96 h post co-transduction with shARID1A and shARID1B (shARID1A/B).

Functional annotation of these 469 epigenetically repressed genes showed a clear enrichment of categories related to invasiveness (Figure 3E, Table S7), including several proteins and regulators of the metastatic process, such as adhesion proteins (i.e. integrins and cadherins) and key regulators of the mesenchymal phenotype (e.g. *LOXL2* and *SNAI2* [5]) (Figure 3C, Table S8). Among these genes, integrins were the most extensively repressed gene family. Integrins are cell-matrix adhesion proteins that act both as sensors and transducers of external stimuli to the cells, and as mechanical effectors of processes like migration and invasion, through their interaction with multiple extracellular matrix (ECM) components such as collagen, fibronectin, or laminin [35]. Among the 26 different human integrin genes, 11 were transcriptionally downregulated after BAF disruption (Figure 3F, Figure S7A-B), 10 of which were assigned at least one chromatin remodelling repressive event in near genomic sites (Figure 3F, Figure S7C), indicating a wide epigenetic repression of this gene family when the structural integrity of the BAF complex is compromised. Interestingly, these BAF-modulated integrins were also modulated after single inhibition of ARID1A or ARID1B, but to a lesser extent (Figure S7D), further highlighting the need for simultaneous inhibition of both proteins for full achievement of BAF inhibition effects. The functional consequences of this epigenetic event were supported by the protein reduction of the two most repressed integrins, namely, ITGB3 and ITGA9 (Figure 3G), both of which have been previously implicated in metastasis [36,37], as well as the clear repressive effect observed on transcriptional signatures related to the activation of integrin signalling (Figure S7E) and protein downregulation of key structural components of downstream pathways such as SRC or ILK, the last of which was transcriptionally and epigenetically repressed (Figure S7F-I).

Additionally, many mesenchymal phenotype-defining genes such as certain cadherins (*CDH2, CDH5, CDH6, CDH11*) or master regulators (*LOXL2, SNAI2*) were also epigenetically repressed upon BAF disruption (Figure 3C). Indeed, the most repressed transcriptional hallmark found in BAF-disrupted cells was epithelial-to-mesenchymal transition (EMT) (Figure 2E, Table S2). The non-epithelial origin of neuroblastoma cells makes them unable to undergo EMT, but recent epigenetic evidence has shown their classification into adrenergic or mesenchymal lineages, with the latter being more undifferentiated, invasive, and drug resistant [5]. Neuroblastoma-specific mesenchymal transcriptional signature was also repressed after BAF disruption in neuroblastoma cells (Figure 3H), involving the epigenetic repression of key regulators of the mesenchymal phenotype, such as *SNAI2, CDH11, CDH2, LOXL2* and *NOTCH2* (Figure 3C and I, Figure S5C).

These results suggest the putative epigenetic control through the BAF chromatin remodelling complex of an invasiveness transcriptional program in neuroblastoma cells, which could be exploited for therapeutic purposes.

### BAF complex disruption represses metastasis-related features of neuroblastoma cells

The drastic modulation of important regulators of the invasiveness and metastatic potential of cancer cells revealed a new oncogenic role of the BAF chromatin remodelling complex in neuroblastoma beyond proliferation. Hence, we decided to validate whether this epigenetic reprogramming manifested itself as a functional phenotypic effect associated with metastasis-related functions.

We first assessed the interaction of neuroblastoma cells with the ECM *in vitro*, specifically with collagen, a ubiquitous integrin ligand and one of the most abundant ECM components [38]. Ablation of the BAF complex resulted in a drastic reduction in the attachment dynamics of neuroblastoma cells to a collagen matrix (Figure 4A), indicating a clear reduction in their affinity for and ability toward adhesion to this ECM component. Accordingly, BAF inhibition reduced the ability of neuroblastoma cells to invade and migrate through a collagen barrier (Figure 4B-C) for a particular period (16 h) at which differences could not be explained by different proliferation rhythms between the experimental conditions. Finally, confocal microscopy of actin filaments by phalloidin staining showed that BAF complex disruption produced a clear change in neuroblastoma cell morphology to a rounded form with the destruction of stress fibres (Figure 4D), cell prolongations that mediate cell motility [39]. Quantification of these prolongations confirmed the observable morphological changes (Figure 4E).

**Fig. 4:**
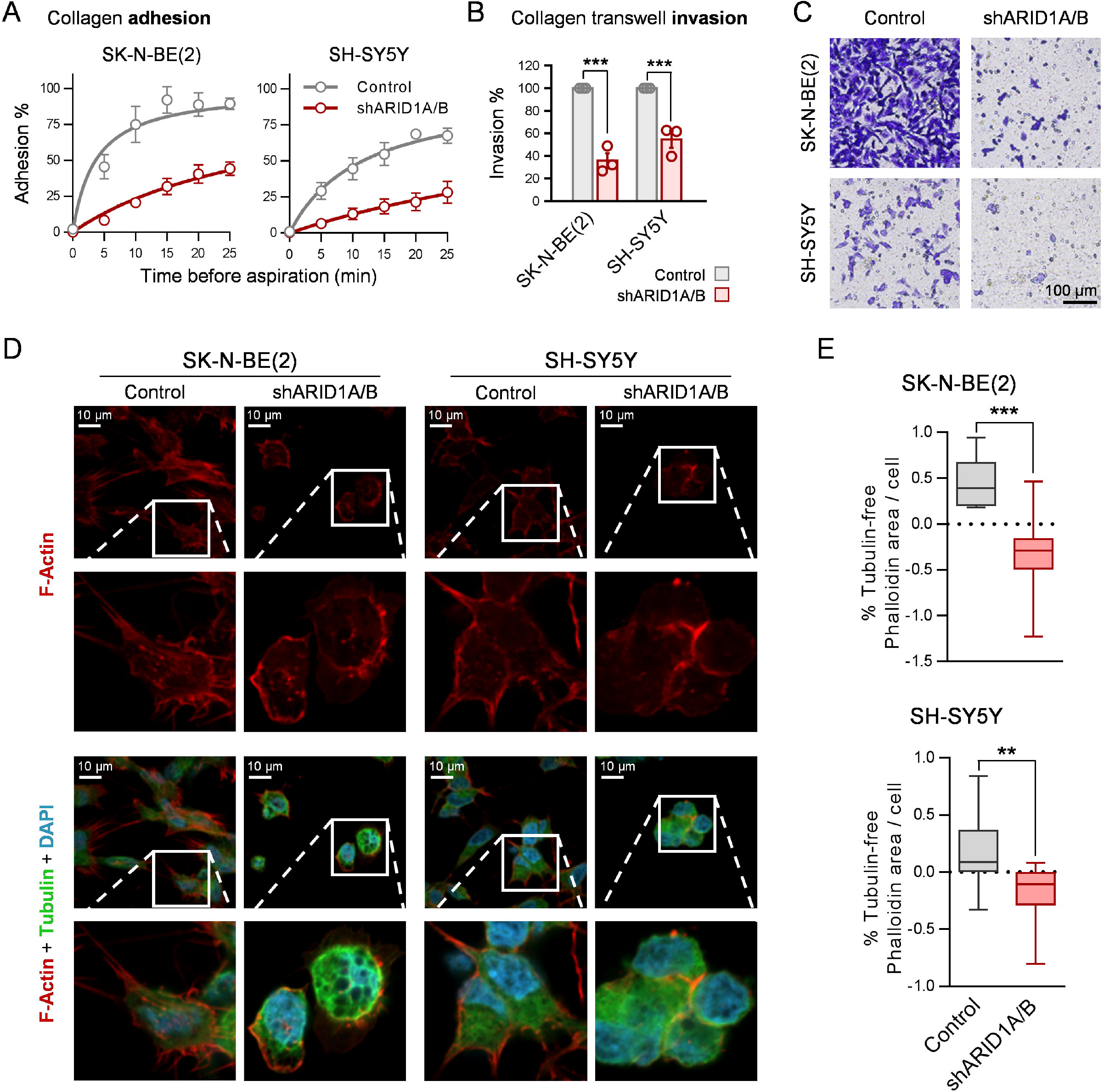
BAF complex disruption reduces collagen adhesion and invasion *in vitro*, and causes morphological changes and destruction of stress fibres in neuroblastoma cells. (**A**) Adhesion assays of neuroblastoma cells to collagen-coated 96-well plates. Cells were aspirated and rinsed at the indicated time points after seeding. Attached cells were quantified using crystal violet staining and absorbance values were normalized to non-aspirated cells. (**B**) Invasion assay with collagen-coated transwells. Neuroblastoma cells (2×10^5^ cells/well) were seeded in collagen-coated transwells and let migrate for 16 h. Invaded cells were quantified by crystal violet staining (**C**) Representative microscopy pictures invaded cells. (**D**) Representative images of neuroblastoma cells stained with phalloidin (F-Actin), anti-tubulin antibody and DAPI by confocal microscopy. (**E**) Quantification of tubulin-free phalloidin area per cell of SK-N-BE(2) (upper graph) and SH-SY5Y (lower graph) cell lines. Ten fields per biological replicate (n = 3) were analyzed. ** means *p* < 0.01; *** means *p* < 0.001.

Altogether, the invasiveness epigenetic program that was deactivated upon BAF disruption exerted deleterious functional consequences *in vitro* on relevant metastasis-related properties in neuroblastoma cells, such as ECM adhesion, motility, and invasiveness.

### Structural integrity of the BAF complex is essential for neuroblastoma metastasis initiation and progression in vivo

Based on *in vitro* evidence, we assessed the effects of BAF structural disruption on the initiation and growth of neuroblastoma metastatic dissemination *in vivo* using mouse models of liver and bone marrow metastasis generated by tail vein injection of SK-N-BE(2) cells [32].

Because the proliferation blockade exerted by BAF disruption on neuroblastoma cells was expected to mask any deleterious effects on their arrival and invasion into the metastasis target organs, we monitored the initial stages of liver colonisation by metastatic cells at single-cell resolution using flow cytometry (Figure 5A). Neuroblastoma cells were detected and counted using the fluorescent mCherry reporter, and their proliferation rate was assessed using a fluorescent proliferation tracing Far Red dye. This double reporter system allowed the discrimination of the human neuroblastoma cell population from mouse hepatocytes (Figure 5B). Only 4 days after injection, a drastic 10-fold reduction in the detection of neuroblastoma cells was already observed in the liver of mice injected with BAF-disrupted (i.e. shARID1A/B) cells, in comparison with those transduced with an empty vector (Figure 5B and C). This decrease was not explained by differences in cell proliferation because the tracer signal was similar between groups at this time point (Figure 5D). At 7 days post-injection, while metastatic control cells had already started populating the liver by proliferation, the few BAF-disrupted cells that had already reached the liver did not increase in number, becoming almost undetectable and impeding proliferation tracer quantification (Figure 5B-D). These results prove that BAF disruption impairs the initial liver colonisation of metastatic cells and not only the posterior metastatic lesion growth through cell cycle arrest.

**Fig. 5:**
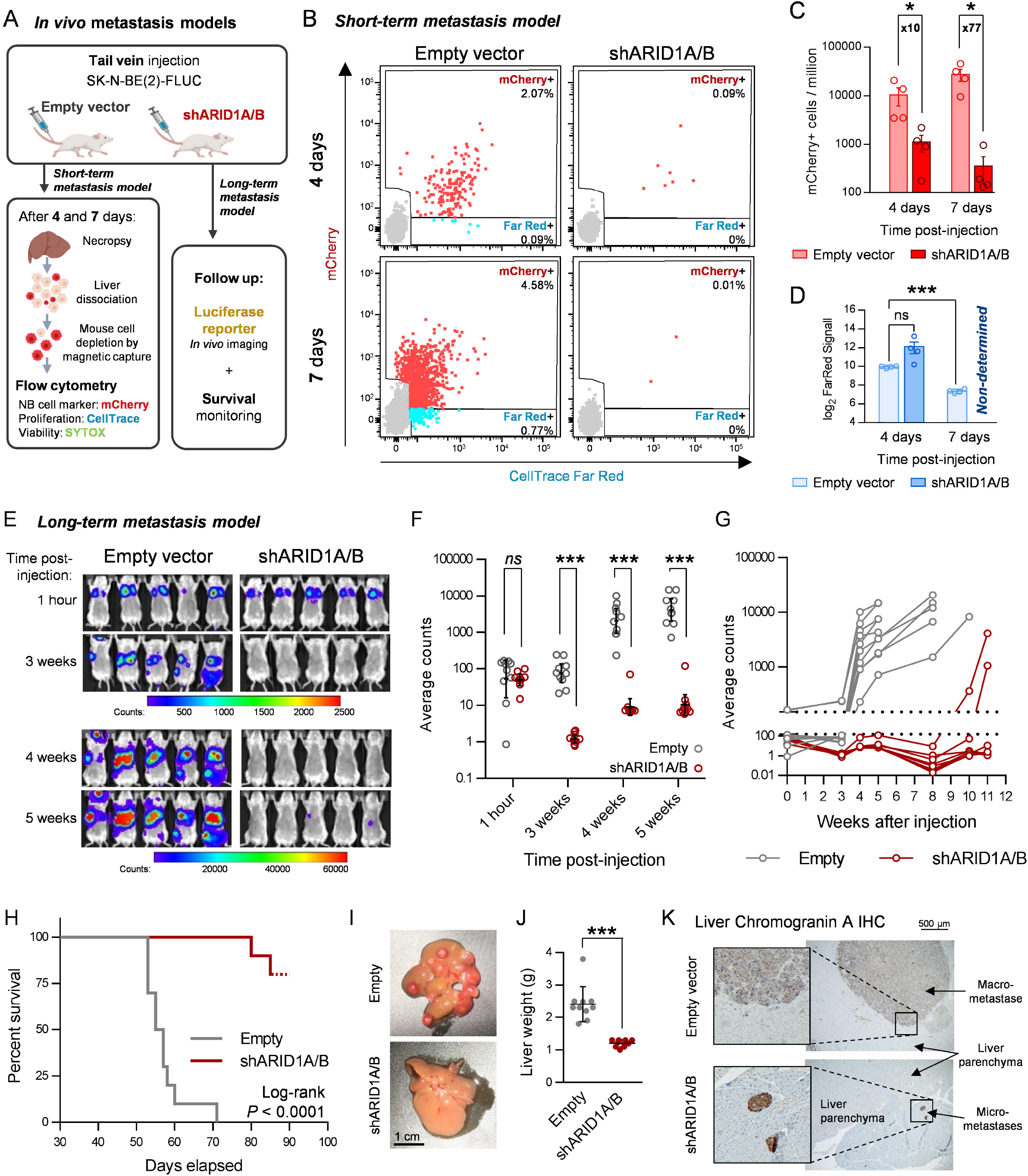
BAF disruption impairs neuroblastoma metastasis initiation and growth *in vivo*. (**A**) Experimental design of the in vivo short- and long-term neuroblastoma metastasis models performed with SK-N-BE(2) cells. (**B**) Representative flow cytometry plots of the short-term metastasis experiment, showing mCherry positive (mCherry+) and FarRed positive/mCherry negative (FarRed+) single living cell populations. (**C**) Quantification of detected mCherry positive cells, expressed in events per million of living single cells (parent gate). Mann-Whitney’s test was performed for statistical comparisons. Fold change between conditions are indicated. (**D**) Average FarRed intensities assessed, when possible, for the mCherry+ population of each experimental group. (**E**) Representative *in vivo* luminescence images of 5 mice per group at the indicated time points. Scale bar represents luminescence counts (photons). (**F**) Luminescence quantification, expressed in average counts, and comparison between experimental groups at the indicated times post-injection. (**G**) Individual mice luminescence quantification follow-up through the entire experiment. (**H**) Kaplan-Meier survival plot comparing mice injected with SK-N-BE(2) transduced with empty vector (Empty) or shARID1A/B. Log-rank test was performed to assess statistical significance. (**I**) Representative images of mouse livers after necropsy. (**J**) Comparison between groups of liver weight at necropsy. (**K**) Representative bright field microscopy images of chromogranin A immunohistochemistry of FFPE liver slides. *ns* means ‘non-significant’; * means *p* < 0.05; ** means *p* < 0.01; *** means *p*< 0.001.

The long-term consequences of this anti-invasive effect were assessed by monitoring metastatic lesion growth and mouse survival, using a luciferase reporter gene for *in vivo* imaging (Figure 5A). Prior *in vitro* analysis revealed that silencing of ARID1A and ARID1B produced a 2.516 ± 0.29 - fold decrease in the luminescence signal (Figure S8A), probably because of metabolic or chromatin accessibility changes. By adjusting with this correction ratio, luminescence *in vivo* imaging at 1-hour post-injection validated the homogeneous injection of viable cells between both experimental groups, with all signals being concentrated in the lungs (Figure 5E-F). *In vivo* weekly imaging showed the disappearance of cells from the lungs, followed by the detection and sustained increase of luminescence signal in multifocal liver and bone marrow metastases in control mice, whereas in animals injected with BAF-disrupted cells, the luciferase signal remained close to background levels and no detectable metastases were observed, except for two mice that developed delayed liver mono-focal metastases (Figure 5E-G, Figure S8B). Although no impact on mouse weight was observed (Figure S8C), BAF disruption extended mouse survival (Figure 5H). While all control mice died between days 53 and 71 post-injection, only 20% of mice injected with BAF-depleted cells developed metastasis-related symptomatology and survived until days 80 and 85 post-injection. Necropsy showed that livers from mice injected with BAF-disrupted cells were devoid of macroscopic lesions, except for two mice that had developed delayed mono-focal metastases, in contrast with the multifocal liver macro-metastases in the empty vector group (Figure 5I, Figure S8D). This effect was further reflected in the liver weight difference between both groups (Figure 5J). Immunohistochemistry analyses identified only one micro-metastasis in slides from three different lesion-free livers from the shARID1A/B group of mice (Figure 5K). Of note, the only two macro-metastases observed in the BAF-disrupted group grew through evasion of the silencing technique, since no downregulation of ARID1A/B, DPF2, or cyclin D1 was observed at the protein level (Figure S6E), reinforcing the impact of BAF complex disruption on metastasis formation and progression.

In summary, BAF structural disruption impairs neuroblastoma metastasis at very early stages by reducing the arrival, invasion, and/or posterior survival of neuroblastoma cells at the metastatic site, an effect later magnified by the strong blockade of cell proliferation. The combination of these effects prevents the formation and growth of macro-metastases and expands the survival of metastatic neuroblastoma models *in vivo*.

## Discussion

Neuroblastoma oncogenic properties reflect aberrant epigenomes [5,6], which require molecular effectors, including chromatin remodellers, to be translated into specific chromatin states. We had already associated the main catalytic subunit of mSWI/SNF remodellers, BRG1, with poor prognostic clinical parameters and oncogenic transcriptional pathways in neuroblastoma [29]. This evidence, together with the recurrent mutations reported for the *ARID1A* and *ARID1B* genes in patient samples [25,26], led us to deepen into the understanding of these epigenetic regulators as plausible therapeutic targets.

Hence, we performed a systematic study of the presence, composition, and biological role of the different mSWI/SNF complexes in neuroblastoma cells. Integral proteomic characterisation of the mSWI/SNF complexes in neuroblastoma cells had not been previously reported. Thus, there was still a need to address their presence and structural integrity to gain a complete understanding of these remodellers in neuroblastoma. The SWI/SNF complex subunits are particularly vulnerable to destabilisation and proteasomal degradation when physically separated from the complex [40]. Therefore, proteomic detection of all known subunits of the three mSWI/SNF subtypes after affinity purification indicated the presence and structural integrity of these subcomplexes in neuroblastoma cells.

ATP hydrolysis is a well-known energy source required for nucleosome mobilisation and chromatin remodelling initiation by the SWI/SNF complexes. However, catalytic inhibition of the complex by degradation of ATPase subunits did not exert observable effects on neuroblastoma proliferation, whereas BAF-specific structural disruption did. Multiprotein complexes concentrate in time and space varied molecular activities for the completion of a multistep process. As such, mSWI/SNF remodellers integrate ATP-dependent DNA translocation, nucleosome anchor, and histone mark binding, among several others, to achieve their functions. Thus, the inhibition of one of its activities could not unveil all the roles of mSWI/SNF, as reported previously with the use of bromodomain inhibitors [41]. Indeed, the SWI/SNF complexes are able to maintain altered unfolded nucleosomes without any catalytic activity [42] and have been found to exert part of their genome-wide functions in an ATPase-independent manner in *Drosophila* [43] and humans [44]. A possible explanation for this phenomenon could be the interplay between the mSWI/SNF complex and transcriptional repressors, such as its antagonist PRC2 [45], conditioning the need for catalytic activity for active chromatin remodelling. Here, we demonstrated a relevant proliferative dependency of neuroblastoma cells on BAF complexes, which is not produced after mere catalytic inhibition but rather requires the full functional depletion of the complex through its structural disruption to be unmasked. In any case, specific full ablation of the BAF complex impaired neuroblastoma proliferation by promoting a strong arrest of the G1 phase and the shutdown of a cell cycle transcriptional signature associated with poor prognosis in patient expression datasets.

Concordantly, BAF disruption exerted a strong protein repression of cyclin D1 and downstream effectors, a regulatory axis crucial particularly for neuroblastoma cells, which have higher levels of dependency for this protein than those of other origins [6]. Our results show that inhibition of this priority neuroblastoma target is one of the potential therapeutic benefits of BAF disruption.

Previous studies have attributed functional consequences to *ARID1A*-loss in the context of *MYCN*-amplified tumours [28], and some authors have postulated *ARID1A* as the main driving tumour suppressor in 1p36-lost *MYCN*-amplified cases [46]. While these studies were focused in the study of one of these subunits alone, we analysed the BAF complete functions by inhibiting both paralogues, *ARID1A* and *ARID1B*, simultaneously, discovering a proliferative dependency on the structural dependency of the complex regardless of *MYCN* status, on the base of the strong cell cycle blockade observed in *MYCN* non-amplified cells (SH-SY5Y). Moreover, the cell cycle-related transcriptional signatures controlled by BAF complex in our experimental setting, described by ARID1A/B combined silencing, although positively correlated with *MYCN* status and expression, is independently associated with poor prognosis of neuroblastoma patients, strongly suggesting that an increased BAF activity has oncogenic consequences, regardless of *MYCN*.

While we previously reported that BRG1 shRNA-silencing impairs neuroblastoma proliferation and triggers apoptosis ([29] and Figure S9A), here we show that BRG1 degradation does not (Figure 1B), a discrepancy attributable to the disruptive effect of shBRG1 on the complex structural integrity (Figure S9B), not produced by ACBI1 (Figure 1C). Intrinsic differences of these loss-of-function strategies, working at pre- and post-translational levels, respectively, may explain this effect. The reason why shBRG1, but not shARID1A/B, triggers apoptosis remains elusive. A possible explanation is that, while shARID1A/B does not alter BRG1 levels (Figure S9C), shBRG1 combines ATPase inhibition and structural disintegration, synergizing and triggering apoptosis. However, ACBI1 synergized with shARID1A/B in cycle arrest, but not in promoting apoptosis (Figure S9D). Potentially, shBRG1 structural effects on other subcomplexes (i.e. PBAF) may be adding an extra layer of stress for neuroblastoma cells, thereby triggering the apoptotic process.

Genome-wide analysis of the specific BAF molecular function, which is promoting chromatin accessibility, has served to narrow down the list of BAF-modulated genes to those directly regulated at the epigenetic level. Notably, ATAC-Seq revealed that the majority of differentially accessible sites underwent repression after BAF disruption, supporting that most of the observed chromatin remodelling events could be attributed to the activity of the BAF complex on these genomic sites. The fact that these events were predominantly located at intergenic and intronic sequences, most probably belonging to distal regulatory elements (i.e. enhancers), is in line with the previously found tendency of BAF complexes towards these sites over promoters [17,34]. Moreover, the neuroblastoma mesenchymal transcriptional signature, repressed after BAF disruption, is determined by a specific super-enhancer configuration [5], and many of these genes (*LOXL2, SNAI2, CDH11, ITGAV*) are associated with chromatin repressive events. This could be a reflection of a putative role of the BAF complex in maintaining open chromatin states at the super-enhancers controlling the expression of this mesenchymal signature.

The merged list of both transcriptionally and epigenetically BAF-modulated genes was markedly enriched with several players related to the cell-matrix interaction, a defining trait of the invasiveness and metastatic potential. Metastatic spread is one of the major challenges in the clinical management of neuroblastoma patients, occurring in 90% of newly diagnosed high-risk cases and in more than 70% of relapsed cases [47,48]. Identifying the molecular mechanisms enabling neuroblastoma cells to undergo every step of the metastatic process is essential for the development of new therapeutic strategies against metastasis formation and progression in these patients. Our findings indicate that the BAF complex allows the expression of an invasiveness epigenomic program involving several adhesion molecules and mesenchymal genes, which is deactivated upon BAF structural disruption. This set of genes includes a wide collection of integrins, some of which have already been reported to have pro-metastatic functions in neuroblastoma, including integrins α1, α3, α4, αv, and β3 [49,50]. Hence, the functional consequences observed in the adhesion, invasion, and metastatic functions of neuroblastoma cells can be partly attributed to the epigenetic repression of this gene family. Either way, we could dissect the detrimental effects on proliferation and prove relevant impairment of neuroblastoma metastatic colonisation caused by BAF disruption in animal models.

While our results clearly show an oncogenic function of the BAF complex in neuroblastoma proliferative and metastatic capacities by controlling poor prognosis-associated transcriptional and epigenetic networks, they apparently conflict with the potential tumour suppressive functions of ARID1A and ARID1B in neuroblastoma, which are supported by different evidences: recurrent deleterious mutations of these two genes [25,26]; the genomic location of *ARID1A* gene in the recurrently deleted 1p36, and its postulation as one of the driving tumour suppressors of this in *MYCN*-amplified neuroblastomas [46]; and functional evidence showing that ARID1A loss promotes adrenergic-to-mesenchymal transition [28] and potentiates invasion [27]. Nevertheless, these studies were based on the single inhibition of one of the two homologous subunits and did not evaluate the effects of combined silencing of ARID1A and ARID1B, which is necessary for full BAF structural disruption. Indeed, proliferative dependency on ARID1B in ARID1A-mutated neuroblastoma cell lines has already been demonstrated in one of these studies [28]. The increased metastatic properties attributed to the loss of one homologue, whether by natural mutations or by experimental loss-of-function studies, could be explained by an overcompensation effect of the remaining one. We hypothesize that the remaining subunit would maintain the structural integrity of the BAF complex and its resulting chromatin states, and would potentiate the expression of an invasiveness epigenomic program, thereby becoming a proliferative and metastatic vulnerability of these cells.

Our work still presents limitations regarding the used models, since although working with different molecular backgrounds (*MYCN*-amplified and *TP53*-deficient SK-N-BE(2) cells; *MYCN*-non-amplified and *TP53*-functional SH-SY5Y) these models neither comprise the totality of neuroblastoma molecular subtypes, nor represent the natural evolution and progression of the disease, as both cell lines were derived from bone-marrow metastases. In order to understand the biological role of BAF complex in the spontaneous process of neuroblastoma metastasis formation, future studies with different neuroblastoma models will definitely be mandatory. Nevertheless, our results do serve to point out a potentially exploitable vulnerability of already-metastatic neuroblastoma cells, which are the main target of interest of anti-metastasis therapies for this disease, since the majority of neuroblastoma metastatic patients debut with already formed metastases [2].

## Conclusions

The specific epigenetic control of such concrete functions by the BAF complex in neuroblastoma cells opens the possibility of reverting an epigenetic program to inhibit several metastatic effectors and regulators simultaneously, with potential as an anti-metastatic epigenetic therapy for neuroblastoma. Furthermore, considering the strong effects of BAF disruption on proliferation through cell cycle arrest and downregulation of one of the most-wanted neuroblastoma therapeutic targets, cyclin D1, these findings reveal BAF complex structural disruption as a potential therapeutic strategy for the treatment of metastatic high-risk neuroblastoma and urge for the design and development of pharmacological structural disruptors of this chromatin remodeller.

## List of Abbreviations

mSWI/SNF: mammalian switch/sucrose non-fermenting
BAF: BRG1-associated factor
PBAF: polybromo-associated BRG1-associated factor
ncBAF: non-canonical BAF
PBS: phosphate-buffered saline
ECM: extracellular matrix
EMT: epithelial-to-mesenchymal transition

## Declarations

### Ethics approval and consent to participate

Not applicable.

### Consent for publication

Not applicable.

### Availability of data and materials

The sequencing datasets generated during the current study are available in the Gene Expression Omnibus (https://www.ncbi.nlm.nih.gov/geo/) under accession numbers GSE202231 (RNA-Seq) and GSE202306 (ATAC-Seq). Neuroblastoma patient sample gene expression analyses were performed with publicly available datasets extracted from R2 genomics analyses and visualization platform (http://r2.amc.nl) under the accession numbers GSE45547 and GSE62564.

### Authors’ contributions

CJ and MFS conceived and designed the study. CJ, RA, MN-R, CS, MM, AM and JR carried out the experiments. CJ, MN-R, LD-J, PL, AS and DL-N analyzed the data. AS, JSdT, DL-N, JR, FP, EdN, SG and LM provided intellectual support for result interpretation and critical revision. CJ and MFS wrote and revised the initial manuscript. All the authors read and approved the final manuscript.

## Acknowledgements

Plasmids bearing Tn5 were a kind gift from Dr. Kim Remans (Protein Core Facility, EMBL) and Dr. Lars Steinmetz (EMBL, Stanford University). FLUC plasmid (CSCW2-Fluc-ImC) was kindly provided by Dr. Rana Moubarak (New York University). We thank all the Protein Expression Core Facility and the Functional Genomics Facility at IRB Barcelona for technical support. We thank the members of the Childhood Cancer and Blood Disorders group (VHIR) for their support. We would like to acknowledge the members of the Laboratory Animal Service and High Technology Unit of VHIR for their technical support. We are grateful to the CNAG-CRG for technical and bioinformatics assistance with transcriptome analyses. We would like to thank Editage (www.editage.com) for English language editing.

## Additional files

*Additional file 1 (.pdf):* Supplementary information. Includes Supplementary Materials and Methods, Supplementary Figures, Supplementary Tables and References.

*Additional file 2 (.xlsx):* RNA-Seq BAF-modulated genes. List of genes transcriptionally modulated in SK-N-BE(2) cells after BAF disruption through ARID1A and ARID1B silencing, ordered by cluster. Log_2_ fold change versus control and adjusted *p*-value are indicated.

*Additional file 3 (.xlsx):* ATAC-Seq and RNA-Seq BAF-downregulated genes. List of genes transcriptionally repressed (RNA-Seq) and with associated chromatin closing events (ATAC-Seq) in SK-N-BE(2) cells after BAF disruption through ARID1A and ARID1B silencing. RNA-Seq expression log_2_ fold change versus control and adjusted *p*-value are indicated.

