## Additional File 1 for "Structural disruption of BAF chromatin remodeller impairs neuroblastoma metastasis by reverting an invasiveness epigenomic program"

Carlos Jiménez *et al*. 2022

Content:


1. **Supplementary Materials and Methods**
2. **Supplementary Figures**
3. **Supplementary Tables**
4. **References**

### 1. **Supplementary Materials and Methods**

*Cell lines and tissue culture*

SH-SY5Y cell line was purchased from American Type Culture Collection (ATCC, Manassas, VA, USA), SK-N-BE(2) cell line was procured from the Public Health England Culture Collection (Salisbury, UK), and COG-N-278 cell line was obtained from the Childhood Cancer Repository (Texas Tech University Health Sciences Center, Lubbock, TX, USA). Neuroblastoma cell lines were cultured, as suggested by Children's Oncology Group Cell Culture and Xenograft Repository, in Iscove's modified Dulbecco's medium (IMDM, Gibco, Amarillo, TX, USA), supplemented with 20% heat-inactivated foetal bovine serum (FBS; South America Premium, Biowest, Nuaillé, France), 1% insulin-transferrin-selenium supplement (Gibco), 100 U/mL penicillin, 100 μg/mL streptomycin (Gibco), and 5 μg/mL plasmocin (InvivoGen, San Diego, CA, USA). The human embryonic kidney cell line HEK293T was purchased from ATCC and cultured in Dulbecco's modified Eagle's medium (DMEM, Gibco), supplemented with 10% heat-inactivated FBS, 100 U/mL penicillin, 100 μg/mL streptomycin, and 5 μg/mL plasmocin. All cultures were maintained at 37 °C in a 5% CO2 saturated atmosphere, and biannually tested for mycoplasma contamination.

*PROTAC degraders*

SMARCA4/SMARCA2 PROTAC degrader ACBI1, and its negative control *cis*ACBI1, was acquired from the public OpnME repository of molecules for research (Boehringer Ingelheim, Ingelheim am Rhein, Germany), resuspended upon arrival in dimethyl sulfoxide (DMSO) at 10 mM, and stored at -20 ºC until use. BRD9 PROTAC degrader (dBRD9) was purchased form Tocris Bioscience (Bristol, UK), resuspended upon arrival in DMSO at 10 mM, and stored at -20 ºC until use.

*Co-immunoprecipitation and Mass Spectrometry*

SK-N-BE(2) and SH-SY5Y cells were grown in three 150 mm dishes for each biological replicate until 80-90% confluence, and subcellular fractionation protocol was performed. Briefly, cells were scraped in cold subcellular fractionation buffer (10 mM PIPES pH 6.8, 300 mM sucrose, 50 mM sodium chloride, 1 mM EDTA, 0.5% Triton X-100 (Sigma-Aldrich, St. Louis, MO, USA)) supplemented with EDTA-free Protease Inhibitors Cocktail, and incubated for 20 minutes on ice before centrifugation at 720 xG for 5 minutes. Supernatants containing cytosolic fractions were discarded and nuclei-containing pellets were resuspended in immunoprecipitation (IP) lysis buffer (50 mM Tris pH 7.5, 300 mM NaCl, 2 mM EDTA, 10% glycerol, 0.1% SDS, 1% Triton X-100) supplemented with protease inhibitors. Nuclei were lysed by incubation on ice for 30 minutes and centrifuged at 13,300 rpm for 20 minutes to discard cell debris. Nuclear lysate supernatant was collected and protein quantified with the same method used for western blot analyses. For each co-immunoprecipitation reaction, 500 μg of nuclear lysates were pre-cleared with 10μL of Protein A-Sepharose beads (Sigma-Aldrich) and 1 μg of Normal Rabbit IgG (Sigma-Aldrich) in IP buffer. Pre-cleared lysates were incubated overnight at a concentration of 1 µg/µL with 5 µg of Rabbit monoclonal anti-BRG1 antibody (ab110641, Abcam, Cambridge, UK) or Normal Rabbit IgG, before addition of 25 µL of Protein A-Sepharose beads and incubation for 2 hours. Beads were then washed three times with 200 mM Ammonium Bicarbonate (ABC; Sigma-Aldrich) and resuspended in 6 M urea (GE Healthcare Life Sciences, Piscataway, NJ, USA) diluted in 200mM ABC. For protein reduction, 10mM dithiothreitol (Sigma-Aldrich) in 200mM ABC was added and incubated for 1 hour at 37ºC with shaking. Iodoacetamide (Sigma-Aldrich) at 20 mM in 200 mM ABC was added for alkylation, and samples were incubated for 30 minutes at room temperature with shaking in darkness. Digestion with 1 µg of sequence-grade trypsin (Promega, Madison, WI, USA) was performed overnight at 37ºC with constant shaking. Beads were pulled-down and samples were acidified with 20 µL of 100% formic acid. For sample desalting, C18 reverse phase UltraMicroSpin columns were used (The Nest Group, Inc., Ipswich, MA, USA). Columns were conditioned with methanol and equilibrated twice with 5% formic acid. Samples were loaded twice into the columns, washed twice with 5% formic acid and eluted with 50% acetonitrile in 5% formic acid before drying using a SpeedVac concentrator (Thermo Scientific, Waltham, MA, USA).

Samples were resuspended and resolved by liquid chromatography prior to analysis by mass spectrometry. Proteomics analyses were performed in the Proteomics Unit of Centre for Genomic Regulation/Pompeu Fabra University (CRG/UPF, Barcelona, Spain). Samples were analysed by LC-MS/MS using a 60-min gradient in an Orbitrap Velos Pro mass spectrometer (Thermo Scientific). As a quality control, BSA controls were digested in parallel and ran between each of the samples to avoid carryover and assess the instrument performance. Samples were searched against SP_Human database, using the search algorithm Mascot v2.6 [1]. Peptides were filtered based on false discovery rate (FDR) and only peptides showing an FDR lower than 5% were retained. Interactome analysis of SMARCA4/BRG1 in SK-N-BE(2) and SH-SY5Y cells was performed using Significance Analysis of INTeractome (SAINT), a software package for scoring protein-protein interactions based on label-free quantitative proteomics data (i.e., spectral counts) in co-immunoprecipitation/mass spectrometry experiments. SAINT allowed to select bona fide interactions and remove nonspecific interactions in an unbiased manner [2].

*Lentiviral transduction*

pLKO.1-puro plasmids carrying shRNA against different genes were purchased from Sigma-Aldrich, Mission shRNA repository (Table S9). Negative controls used were pLKO-Non-Silencing Control, expressing a non-targeting shRNA, or pLKO-empty, not expressing any shRNA, acquired from AddGene repository (Plasmid #8453; Watertown, MA, USA). Lentiviruses were generated in HEK293T cells using previously described methods [3,4]. HEK293T cells were transfected with a mixture of the specific shRNA plasmid together with lentiviral envelope and packaging vectors pMD2G and psPAX2 with Lipofectamine 2000 (Invitrogen, Waltham, MA, USA) following manufacturer's instructions. Two days after transfection, lentiviral supernatant was collected, centrifuged for 5 minutes at 1500 rpm to discard floating HEK293T cells, and passed through a 0.45 µm syringe filter (Fisher Scientific, Waltham, MA, USA). Lentiviral supernatants were frozen at -80ºC and stored until further use. Transduction of neuroblastoma cell lines with lentiviral particles was performed by seeding $5 \times 10^5$ neuroblastoma cells in 60 mm plates with lentiviral supernatant diluted in IMDM 20% FBS medium at different dilutions, depending on the cell line. The next day after seeding and transduction, lentiviral supernatant was replaced by fresh medium. Transduced cells were maintained in culture, and used at the indicated time points. When performing combination transduction experiments, double concentration of control lentiviral supernatant was used to compensate viral doses, and in the case of single inhibition in these experiments, control lentiviral supernatant was added to the specific shRNA of each condition to equate the amount of lentiviral supernatant in all the samples.

*Western Blot*

From *in vitro* experiments, cells were harvested, pelleted and lysed by resuspension in 5-10 times their volume of RIPA lysis buffer (Thermo Scientific) supplemented with EDTA-free Protease Inhibitor Cocktail (Roche, Basel, Switzerland) and Phosphatase Inhibitor Cocktails 2 and 3 (Sigma-Aldrich). For nuclear extract preparation, cells were resuspended in subcellular fractionation buffer (described in *Co-immunoprecipitation* section), incubated for 20 minutes on ice before centrifugation at 720 xG for 5 minutes, and nuclei lysed with RIPA buffer. In the case of frozen tissues, pieces of 2 to 5 mm diameter were immersed in 200 to 500 µL of the same RIPA buffer, and each fragment was disrupted with 20 ceramic beads with a Bead Ruptor 12 (Omni, Kennesaw, GA, USA), using one cycle of agitation of 20 seconds at a speed of 5 m/s. In both cases, samples were incubated for 20 minutes on ice to allow cell lysis and cleared by centrifugation at maximum speed (i.e. 13,300 rpm) at 4ºC for 15 minutes. Supernatant fraction was kept and protein concentration quantified with DC protein assay (Bio-Rad, Hercules, CA, USA). A total of 30 µg of each protein sample were heated at 70ºC for 10 minutes, resolved onto precasted NuPAGE 4-12% Bis-Tris polyacrylamide gels (Invitrogen) and posteriorly transferred onto polyvinylidene fluoride (PVDF) membranes (GE Healthcare Life Sciences). Membranes were blocked for 1 hour in 5% Albumin Bovine Fraction V (BSA; NZYTech, Lisbon, Portugal) or 5% non-fat dried milk (PanReac AppliChem ITW Reagents, Castellar del Vallès, Spain) diluted in tris-buffered saline 0.1% Tween 20 (Sigma-Aldrich) (TBS-T) and incubated overnight incubation with primary antibodies diluted in 5% BSA or 5% milk TBS-T (Table S10). Membranes were then incubated for 1 hour with horseradish peroxidase (HRP)-conjugated secondary antibodies and developed with Enhanced Chemiluminescence (ECL) (GE Healthcare Life Sciences).

*siRNA transfection*

Sets of custom siRNA duplexes against each targeted gene (i.e. ARID1A and ARID1B) with [dT][dT] overhangs were purchased from Sigma-Aldrich, based on validated target sequences extracted from literature (Table S9). Block-iT fluorescent siRNA control (Invitrogen) was used as negative control and to monitor transfection efficiency. Neuroblastoma cells (2.5 × $10^5$ cells/mL) were transfected with 25 nM of the indicated siRNAs with Lipofectamine 2000, following the manufacturer's instructions. When performing transfection experiments combining different siRNAs, 25 nM siRNA was used for each gene to maintain silencing performance, rising total siRNA concentration to 50 nM. Therefore, concentration of the negative control siRNA was doubled to 50 nM, to compensate doses. In the case of single inhibition in these experiments, negative control siRNA was added to the specific siRNA of each condition to equate siRNA concentrations to 50 nM in all the conditions. Proliferation assays with siRNA-transfected cells were performed by trypan blue exclusion assay using the Cell Counter EVE (NanoEntek, Seoul, South Korea).

*RNA-Sequencing (extended)*

RNA was extracted from 72 hours-transduced 2.5 × $10^5$ SK-N-BE(2) in 6-well plates in biological triplicates by scraping in Qiazol lysis buffer (Qiagen, Hilden, Germany) and purification with miRNeasy mini extraction kit (Qiagen), following the manufacturer's instructions, with an additional in-column step of DNAse I treatment (Qiagen). Total RNA was eluted in 20 µL of nuclease-free water, fluorescently quantified using Qubit RNA HS Assay (Invitrogen) and quality checked by analysis with RNA 6000 Nano Assay on a Bioanalyzer 2100 (Agilent, Santa Clara, CA, USA). All samples contained enough material (> 1 µg) of high RNA quality (RIN 10/10). Library preparation and sequencing was performed at the National Center of Genomic Analyses (CNAG-CRG, Barcelona, Spain). The RNA-Seq libraries were prepared following the TruSeq Stranded mRNA LT Sample Prep Kit protocol (Illumina, San Diego, CA, USA) and sequenced on NovaSeq 6000 (Illumina).

RNA-Seq reads were mapped against human reference genome (GRCh38) using STAR software version 2.5.3a [5] with ENCODE parameters. Genes were quantified using RSEM version 1.3.0 [6] with default parameters and annotation file from GENCODE version 34. Differential expression analysis was performed with DESeq2 v1.26.0 R package [7] using a Wald test to compare control and problem samples. Differentially expressed genes were those with *p*-value adjusted < 0.05 and absolute fold-change (FC) > 1.5, or more restrained thresholds, when indicated. Functional enrichment analysis of Hallmarks gene set collection from MSigDB database were performed using Gene Set Enrichment Analysis (GSEA) software [8,9]. Heatmaps were generated by normalizing the normalized counts of each gene by the average counts of the gene in all conditions, and $\log_2$ transformation. This value was represented in a colour gradient heatmap using Microsoft Office Excel (Microsoft Corporation, Redmond, WA, USA) and TM4's Multiple Experiment-Viewer MeV [10] softwares.

*Cell cycle analysis*

Cell cycle analysis was performed by the propodium iodide method. SK-N-BE(2) and SH-SY5Y transduced cells were fixed 96 h after transduction in 70% ice-cold ethanol overnight at -20°C, at a density of $10^6$ cells/mL. Fixed cells were washed twice with PBS and resuspended in a staining solution containing 15 μg/mL propidium iodide (Sigma-Aldrich), 1.14 mM sodium citrate (Sigma-Aldrich), and 0.3 mg/mL RNase A (PanReac AppliChem ITW Reagents) in PBS at of $10^6$ cells/mL. Cells were incubated at room temperature (20–25 °C) in the staining solution for at least 30 minutes prior to FACSCalibur flow cytometer analysis (BD Biosciences, Franklin Lakes, NJ, USA). Flow cytometry results were analysed using the FlowJo v10.8 Software (BD Biosciences).

*Cell death assays*

Apoptotic cell death was analysed by staining chromatin with Hoechst 33342 (Sigma-Aldrich). Hoechst staining assays were performed on neuroblastoma living cells plated in 24-well plates ($8 \times 10^4$ cells/well). Twenty-four hours after plating, cells were stained with 0.05 mg/ml Hoechst for 30 minutes at room temperature. Stained nuclei were observed and photographed under ultraviolet fluorescence microscopy.

CellTox Green Cytotoxicity Assay (Promega) kit was used for the determination of cell toxicity involving permeabilization of cell membrane and releasing of free genomic DNA. Transduced neuroblastoma cells were seeded on black opaque 96-well plates ($2 \times 10^4$ cells/well). Twenty-four hours later, the CellTox reaction was performed on these same plates by addition of the fluorescent dye, following the manufacturer's protocol. Cell lysis solution included in the kit was used as a positive technical control of cell death. Fluorescence was measured using an Appliskan (Thermo Scientific) microplate reader. Fluorescence signal was normalized against each control.

*ATAC-Sequencing (extended)*

ATAC-Seq samples were processed simultaneously in triplicates as previously reported [11], with minor modifications. A total of $5 \times 10^5$ SK-N-BE(2) cells were obtained by trypsin digestion, washed with cold DPBS (Gibco) and lysed in fresh lysis buffer from [11]. For tagmentation, Tn5 E54K, L372P was expressed and purified from pETM11 vector as described in [12]. Tn5 (0.35 mg/mL) loading was done with linker oligonucleotides (Tn5ME-A/Tn5MErev and Tn5ME-B/Tn5MErev) as previously reported [12] for 60 minutes at 23ºC. Loaded Tn5 was purified with a 30 KDa Amicon Ultra 0.5 centrifugal unit (Sigma-Aldrich) column and diluted to a final concentration of 0.1 mg/ml with glycerol 25%. Tagmentation reaction was performed in a 50 μL reaction (25 uL tagmentation mix, 5 μL of nuclei ($10^5$ cells/uL), 5 μL of loaded Tn5 and, 15 μL of nuclease free water) for 60 minutes at 37 ºC followed by 5 mins at 80 ºC for heat inactivation. Immediately after samples were purified using MiniElute PCR Purification kit (Qiagen). The entire Purified DNA (10 μL) was amplified in a 50 μL reaction with NEBNext 2X Ultra Mastermix (New England Biolab, Ipswich, MA, USA) with combinatorial dual index primers i5 and i7 primers [12] for 12 cycles. HPLC purified indexing primers were obtained from Integrated DNA Technologies (IDT, Newark, NJ, USA). After PCR amplification, samples were purified by Ampure Beads 0.9X ratio (Beckman Coulter, Brea, CA, USA) following manufacturer's instructions. DNA was eluted with 20 µL of Elution Buffer (Qiagen) and libraries were migrated with an Agilent High Sensitivity DNA and Agilent 2100 instrument following manufacturer's instructions (Agilent Technologies). Samples were pooled and sequenced in a NovaSeq 6000 flow cell with paired-end 100 cycles.

ATAC-Seq analyses were performed with nf-core ATAC-seq pipeline v.1.2.1 [13] with default parameters except for the human reference genome (GRCh38.104), modified to include the sequences of the shRNAs used in each of the experimental conditions reference genome (Table S9). Accessibility peaks of representative genomic regions were obtained with the Integrated Genome Viewer [14]. Differentially accessible peaks were determined by DESeq2 by comparing Non-Silencing Control to shRNA combinations (FDR < 0.01). Genes associated with differentially accessible peaks, by means of genomic proximity, were overlapped with differentially expressed genes obtained by RNA-Seq. Density plot and heatmap occupancy were obtained from the alignment files with plotHeatmap and plotProfile functions of the deepTools pipeline [15], using Galaxy platform [16], over the peaks of interest.

*Immunofluorescence*

Actin filaments were stained with the Phalloidin dye. A total of $2 \times 10^5$ cells per well were seeded in collagen-coated glass cover slips in 24-well plates and grown for 2 days. Next, cells were rinsed twice with PBS and fixed with 4% paraformaldehyde for 10 minutes at room temperature. Cells were washed three more times in PBS and incubated in glycine 0.1 M in PBS at room temperature for 5 minutes under soft agitation. After 2 more washes with PBS, cells were permeabilized with 0.1% Triton X-100 in PBS at room temperature. Two more washes with PBS were performed prior blocking with 3% BSA in PBS for 60 minutes at room temperature under soft agitation. After one more wash with PBS, cover slips were incubated with the staining solution containing phalloidin-iFluor 594 (Abcam) diluted according manufacturer's instructions, monoclonal Anti-β-Tubulin−FITC (Sigma-Aldrich) 1000-times diluted and DAPI 10 µ/mL (Invitrogen) in 3% BSA in PBS, for 1 hour at room temperature under soft agitation, in a dark wet chamber. After 3 final washes with PBS, cover slips were mounted onto microscopy slides using ProLong Diamond Antifade Mountant (Invitrogen), and visualized with a ZEISS LSM 980 confocal microscope (Oberkochen, Germany). Ten random fields were acquired for each biological replicate and processed using ImageJ software. Number of cells per field was counted using DAPI staining of nuclei, and area stained with phalloidin and anti-tubulin was calculated. Tubulin-positive area percentage was used as a reference of the surface occupied by the main body of the cell, whereas area percentage of actin filaments protruding from this main body was considered as a quantification measure of stress-fiber protrusions. Thus, quantification of filamentous actin protrusions for each field was performed by calculating the percentage of tubulin-free phalloidin area per cell, by subtracting the tubulin area percentage to the phalloidin area percentage.

*In vitro luciferase assay*

Dual-Glo Luciferase Assay System (Promega) was used to measure the luciferase signal of neuroblastoma cells *in vitro*. Different numbers of FLUC-transduced SK-N-BE(2) cells (1.25, 2.5 and $5 \times 10^5$ per well) 120 hours after transduction were plated in white opaque 96-well plates. Cell lysis and luciferase reactions were performed following the manufacturer's recommendations. Luminescence was measured using an Appliskan microplate reader. Luciferase signal was linearly correlated to the number of viable cells, confirming the reproducibility and feasibility of the technique and the results.

*Immunohistochemistry*

Mice livers were fixed in 10% formalin solution (Sigma-Aldrich) overnight, and washed twice with PBS before embedded in paraffin and sliced. Tissue sections were deparaffinized overnight at 60 °C and rehydrated using graded alcohols. Heat-induced antigen retrieval was performed using citrate buffer (pH 6, 4 min, 11 5°C) in a pressurized heating chamber. Chromogranin A primary antibody (Roche 760-2519, diluted 1:20) was incubated overnight at 4°C after blocking endogenous peroxidase. Tissue sections were incubated with secondary antibody (Dako) for 30 min at room temperature, developed using diaminobenzidine (Dako), and counterstained using hematoxylin.

*Neuroblastoma patient sample dataset analyses*

K-means clustering module of the R2 genomics analyses and visualization platform (department of Oncogenomics, Academic Medical Center of the University of Amsterdam; http://r2.amc.nl) was used to generate two predictive gene-signatures, based on genes downregulated upon BAF inhibition. Two independent neuroblastoma patient cohorts were used for the study: SEQC (n = 498; GSE62564) [17] and Kocak (n = 649; GSE45547) [18]. To select gene-signatures, data obtained from RNA-Seq and ATAC-Seq analysis was used. For the first gene-signature, genes significantly downregulated in RNA-Seq data ($log_2FC < -1.5$ and adjusted $p$-value $< 0.001$) in response to BAF depletion and included in the repressed cell cycle-related hallmarks (GSEA) were selected. For the second gene-signature, genes repressed in RNA-Seq data ($log_2FC < -0.75$ and adjusted $p$-value $< 0.05$), with associated chromatin repressive events assessed by ATAC-Seq, and included in the repressed cell cycle-related hallmarks were selected. Limited by the availability of probes in both datasets, for k-means analysis the first gene-signature included 171 genes, and the second one included 26 genes. In both cases the number of draws was set 10x10. Heatmaps represent z-score normalized expression. Kaplan Meier, based on the clusters, were generated using overall survival and the log-rank test was performed to assess differences between groups. Univariate and multivariate Cox proportional hazard regression analyses were used to assess the prognostic significance of BAF score on overall survival. These statistical analyses were performed using the IBM SPSS 21 software.

In addition, based on the gene-signatures, a z-score was calculated for each patient (by using the R2 Heatmap module) to study the correlations with clinical data (INSS and Risk Group).

*Statistical analyses*

Unless otherwise stated, graphs represent the average of three independents replicates, and standard error of the mean (S.E.M.) is represented by error bars. Statistical significance was determined using GraphPad Prism 6 Software, using two-tailed Student's *t*-test for comparisons between two conditions and one or two way-ANOVA followed by Sidak's test for multiple comparisons, unless otherwise stated.

2. **Supplementary Figures**

Figure S1:

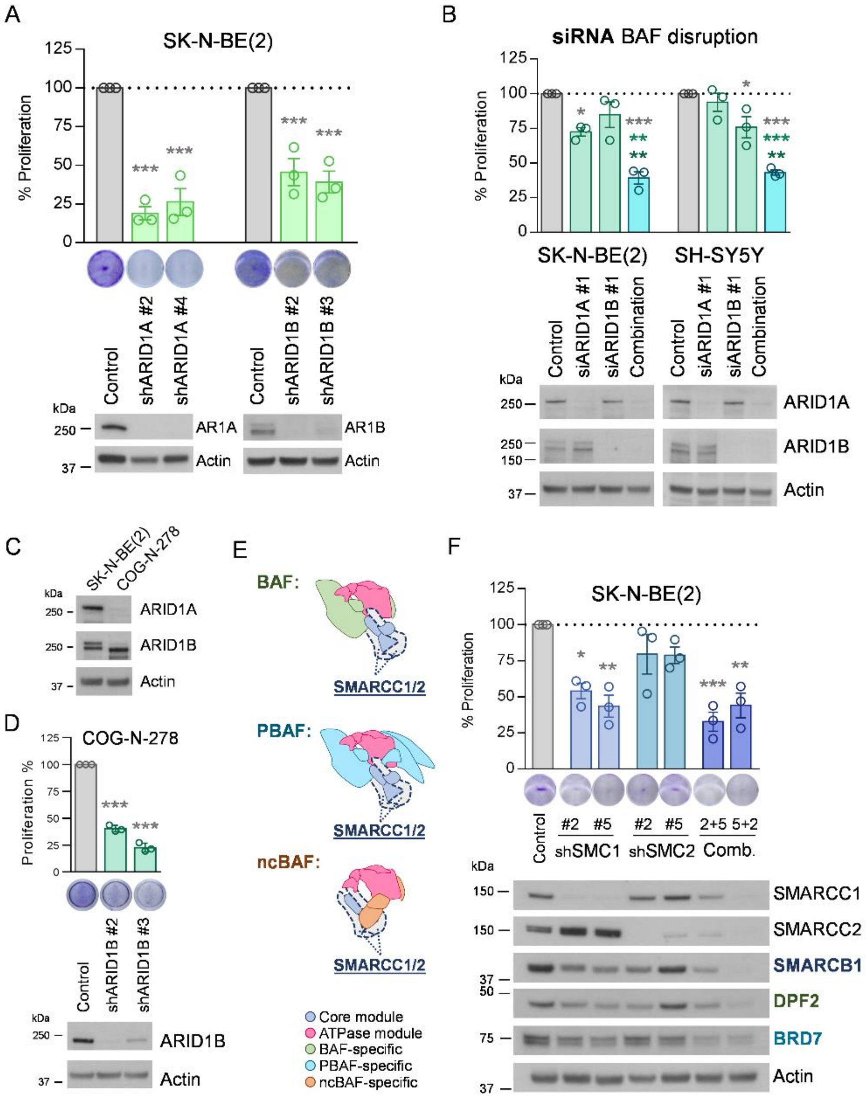


**Fig. S1**: Validation of the effects of BAF complex structural disruption on neuroblastoma proliferation. (**A**) Knockdown of ARID1A and ARID1B with two different shRNA lentiviral vectors for each protein in SK-N-BE(2) cells. Upper panel, proliferation assays of cells seeded in 24-well plates 72 h after transduction and grown for 96 h. Lower panel, knockdown validation by western blot of the target proteins 96 h post-transduction. (**B**) Knockdown of ARID1A and ARID1B separately or in combination using siRNAs in SK-N-BE(2) and SH-SY5Y cells. Upper panel, proliferation analysis of cells transfected in 60-mm plates and grown for one week before trypsinizing and scoring the number of viable cells using the trypan blue exclusion assay. Lower panel, western blot of the indicated proteins 96 h post-transfection validating protein knockdown. (**C**) Western blot analysis of ARID1A and ARID1B in SK-N-BE(2) and COG-N-278 cells. (**D**) Upper graph, proliferation assay of COG-N-278 cells seeded 72 h post-transduction with shARID1B and grown for 96 h. Lower, western blot validation of ARID1B knockdown in 96 h shARID1B transduced COG-N-278 cells. (**E**) Schematic representation of the full mSWI/SNF disruption strategy, showing the key core subunits selected for gene silencing. (**F**) Knockdown of SMARCC1 (shSMC1) and SMARCC2 (shSMC2) with two different shRNA lentiviral vectors for each gene, and two combinations of shRNAs for both proteins, in SK-N-BE(2) cells. Upper panel, proliferation assay of cells seeded 72 h post-transduction in 24-well plates and grown for 96 h. Lower, western blot analysis of the target proteins and other mSWI/SNF subunits 96 h after transduction. * means P < 0.05; ** means P < 0.01; *** means P < 0.001.

Figure S2:

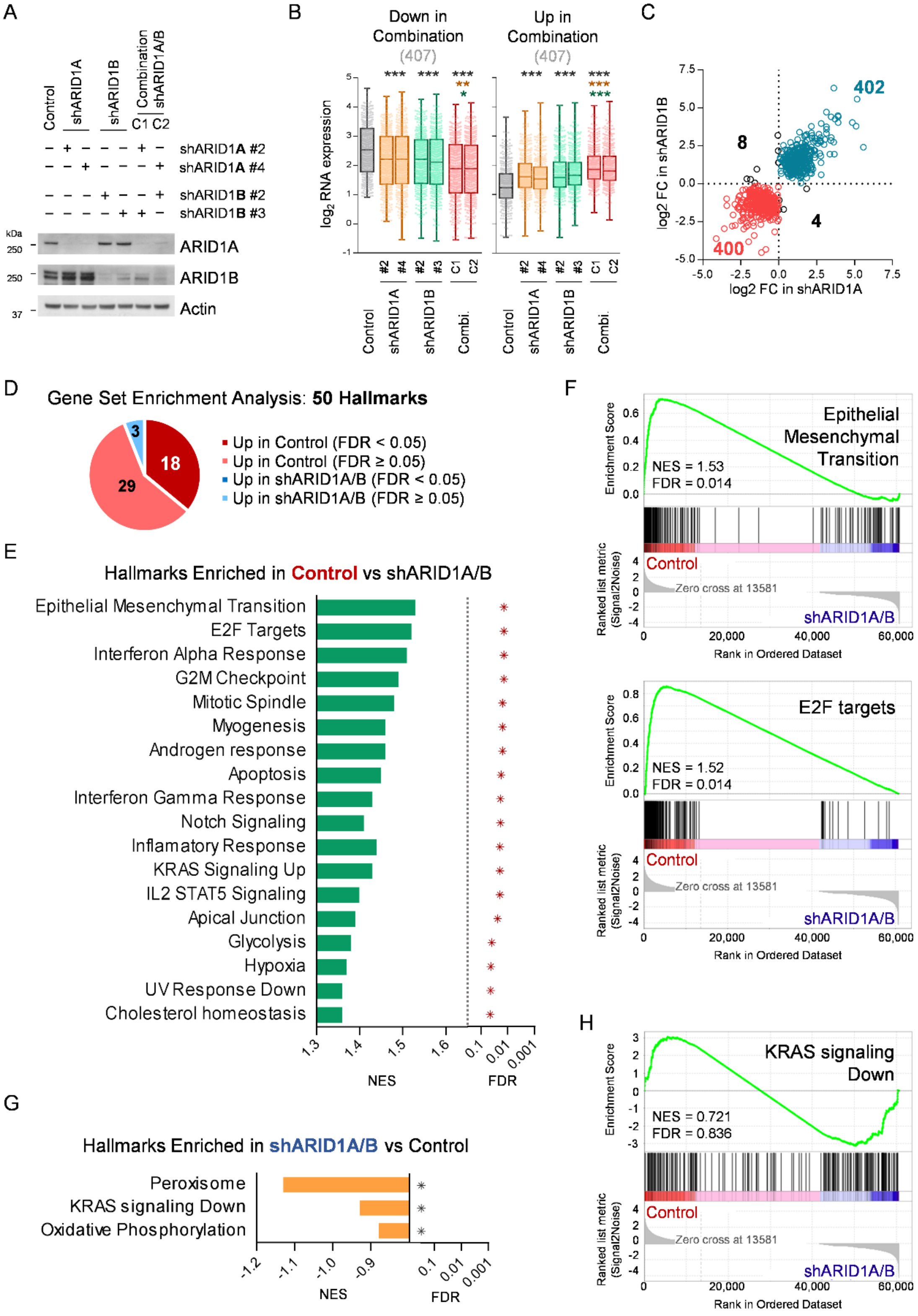
A
Control
shARID1A
shARID1B
Combination shARID1A/B
C1 C2
shARID1A #2
shARID1A #4
shARID1B #2
shARID1B #3
kDa
250
ARID1A
250
ARID1B
37
Actin
B
Down in Combination
(407)
Up in Combination
(407)
log2 RNA expression
Control
shARID1A
shARID1B
Combi.
C
402
8
4
400
log2 FC in shARID1B
log2 FC in shARID1A
D
Gene Set Enrichment Analysis: 50 Hallmarks
3
18
29
Up in Control (FDR < 0.05)
Up in Control (FDR ≥ 0.05)
Up in shARID1A/B (FDR < 0.05)
Up in shARID1A/B (FDR ≥ 0.05)
E
Hallmarks Enriched in Control vs shARID1A/B
Epithelial Mesenchymal Transition
E2F Targets
Interferon Alpha Response
G2M Checkpoint
Mitotic Spindle
Myogenesis
Androgen response
Apoptosis
Interferon Gamma Response
Notch Signaling
Inflamatory Response
KRAS Signaling Up
IL2 STAT5 Signaling
Apical Junction
Glycolysis
Hypoxia
UV Response Down
Cholesterol homeostasis
NES
FDR
F
Epithelial Mesenchymal Transition
NES = 1.53
FDR = 0.014
Enrichment Score
Control
Zero cross at 13581
shARID1A/B
Ranked list metric (Signal2Noise)
Rank in Ordered Dataset
E2F targets
NES = 1.52
FDR = 0.014
G
Hallmarks Enriched in shARID1A/B vs Control
Peroxisome
KRAS signaling Down
Oxidative Phosphorylation
NES
FDR
H
KRAS signaling Down
NES = 0.721
FDR = 0.836

**Fig. S2**: Transcriptome effects of BAF complex structural disruption. (**A**) Western blot validation of ARID1A and ARID1B single or combined knockdowns in SK-N-BE(2) cells at 72 h after transduction, analyzed in parallel to samples used for RNA-Seq analysis. (**B**) RNA expression comparison of 814 BAF-modulated transcripts, split in down- or up-regulated, among experimental groups. Graph represents the $log_2$ transformed RNA-Seq normalized counts for each gene. (**C**) Scatter plot comparing the expression fold change (FC) with respect to control of the 814 BAF-modulated transcripts between single ARID1A and ARID1B inhibitions. (**D**) Pie chart review of gene set enrichment analysis of 50 hallmarks from MSigSB on RNA-Seq expression data from SK-N-BE(2) comparing control against combination of shARID1A and shARID1B (shARID1A/B). (**E**) Normalized Enrichment Score (NES) and False Discovery Rate (FDR) of the 13 gene sets significantly enriched in control. (**F**) Enrichment plots of two of the gene sets most enriched in the control versus combined inhibition of ARID1A and ARID1B, Epithelial Mesenchymal Transition and E2F targets, consisting of 200 genes each one. (**G**) NES and FDR of the 3 gene sets in enriched in shARID1A/B cells versus control. (**H**) Example enrichment plot of one of the non-enriched gene sets, KRAS Signaling Down, consisting of 200 genes.

Figure S3:

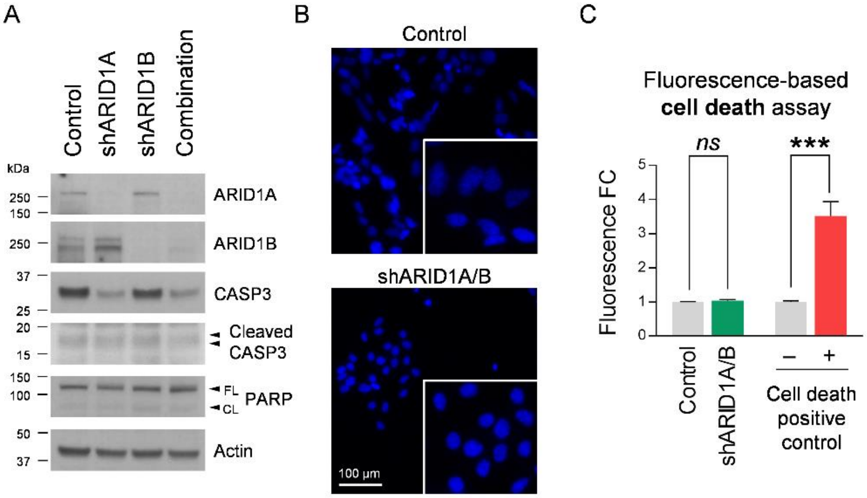


**Fig. S3**: BAF complex disruption does not trigger cell death in SK-N-BE(2) neuroblastoma cells. (**A**) Western blot analysis of the full length (FL) and cleaved (CL) forms of caspase 3 (CASP3) and PARP, 10 days after transduction with shARID1A/B. (**B**) Representative fluorescence images of Hoechst stained SK-N-BE(2) cells at the same time point. (**C**) Fold change (FC) fluorescence based cell toxicity assay of SK-N-BE(2) at the same time point, using CellTox kit (Promega). Lysis solution from the kit was used as cell death positive control. *ns* means ‘non-significant’; *** means $P < 0.001$.

Figure S4:

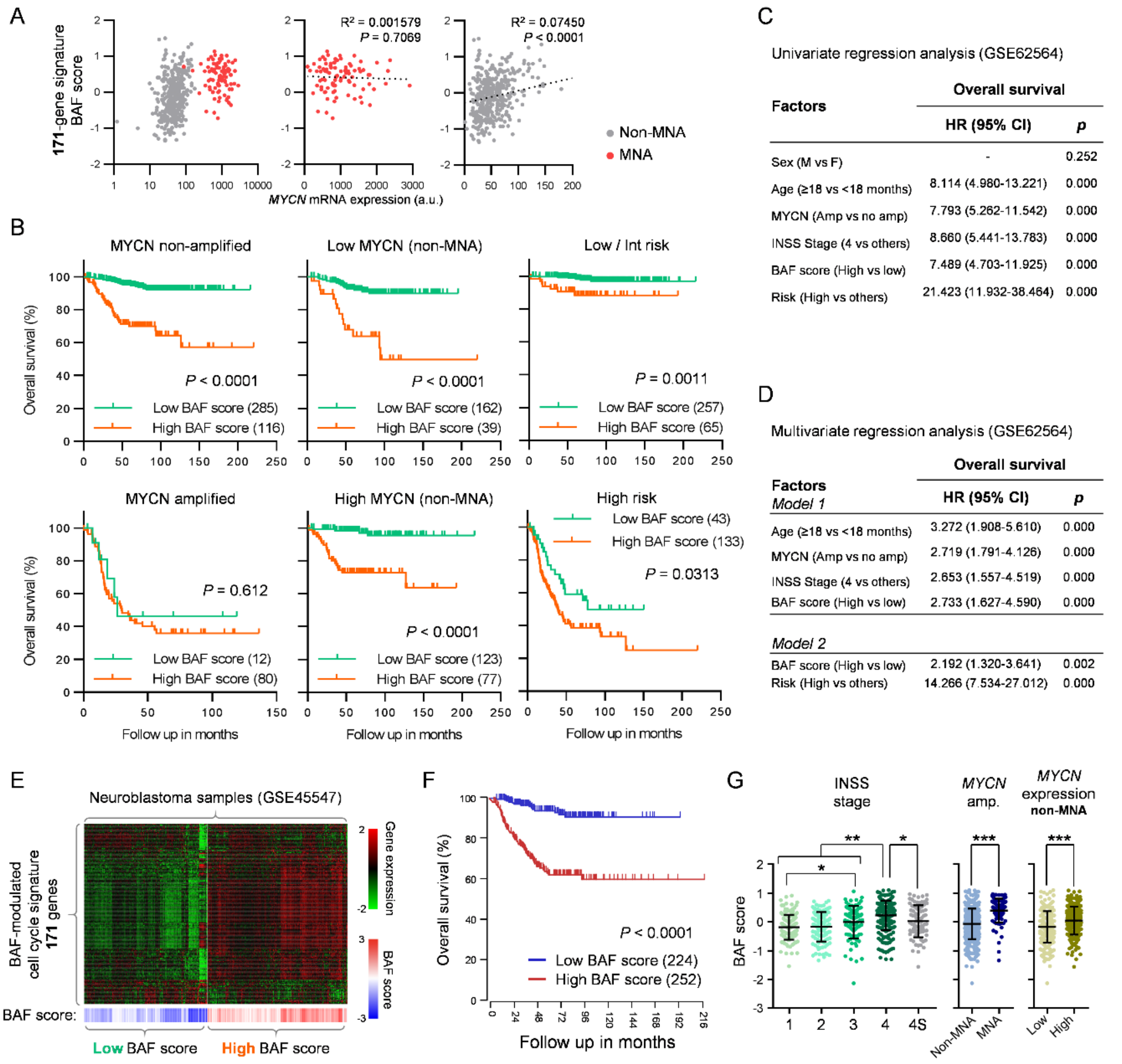


**Fig. S4**: BAF-modulated cell cycle transcriptional signature has independent prognostic value in neuroblastoma. (**A**) Regression analyses between *MYCN* mRNA expression and BAF cell cycle-related signature (171 genes) score in GSE62564 dataset, separating groups according to *MYCN* amplification (MNA). (**B**) Kaplan-Meier plots comparing the overall survival between patients (GSE62564) with high or low BAF score, in separated groups according to MYCN amplification (MNA), *MYCN* mRNA expression levels (above or below median) in non-MNA cases, and risk stratification. (**C**) Cox univariate regression analysis of overall survival with different clinic-pathological features. (**D**) Cox multivariate regression analysis of overall survival confirms BAF signature score as an independent prognostic marker in neuroblastoma. (**E**) An independent cohort of 649 patients (GSE45547) was used to validate the prognostic value of BAF cell cycle signature. Heatmap shows mRNA expression of 171 genes from BAF signature, with patients unbiasedly clustered into high and low BAF score groups. (**F**) Kaplan-Meier plots comparing the overall survival of high and low BAF score group of patients from GSE45547. (**G**) Comparison of BAF signature score between INSS stages, *MYCN* amplification (MNA), and *MYCN* mRNA expression levels (above or below median) in non-MNA cases (GSE45547). * means P < 0.05; ** means P < 0.01; *** means P < 0.001.

Figure S5:

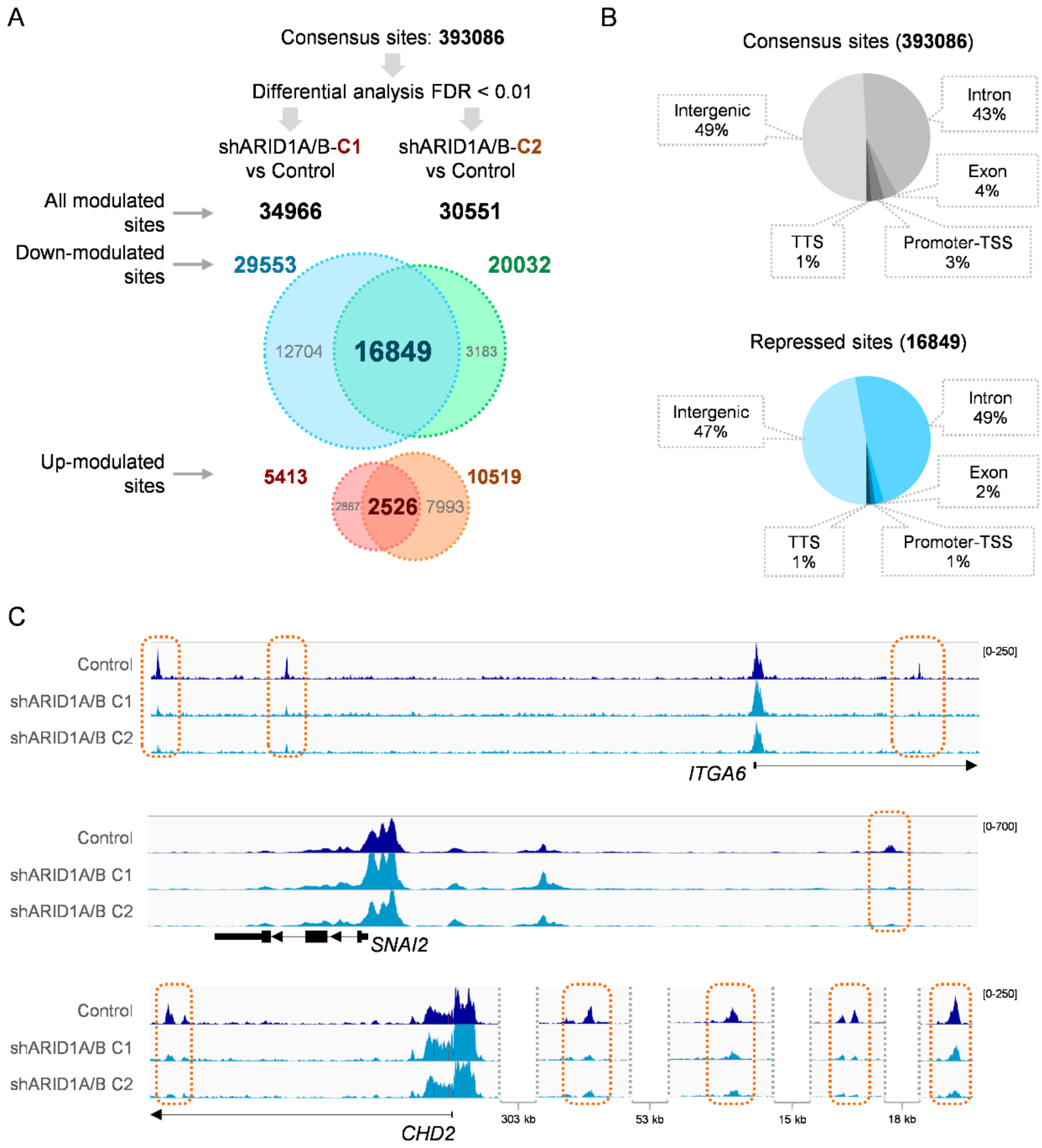


**Fig. S5**: Chromatin accessibility repression after BAF complex disruption. (**A**) ATAC-Seq results analysis workflow for the identification of chromatin accessibility repressive events in SK-N-BE(2) cells after BAF disruption. Those peaks significantly and consistently modulated in both combinations of shRNAs against ARID1A and ARID1B were selected. (**B**) Genomic annotation proportions of all detected accessible sites and of only repressed sites after BAF disruption. (**C**) Representative genome browser ATAC-Seq coverage images of repressed sites after BAF disruption in genes also repressed at the mRNA level, *ITGA6*, *SNAI2* and *CHD2*.

Figure S6:

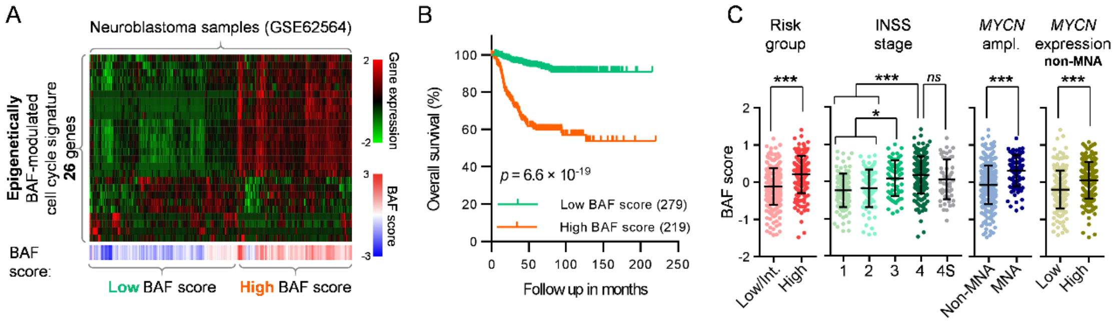


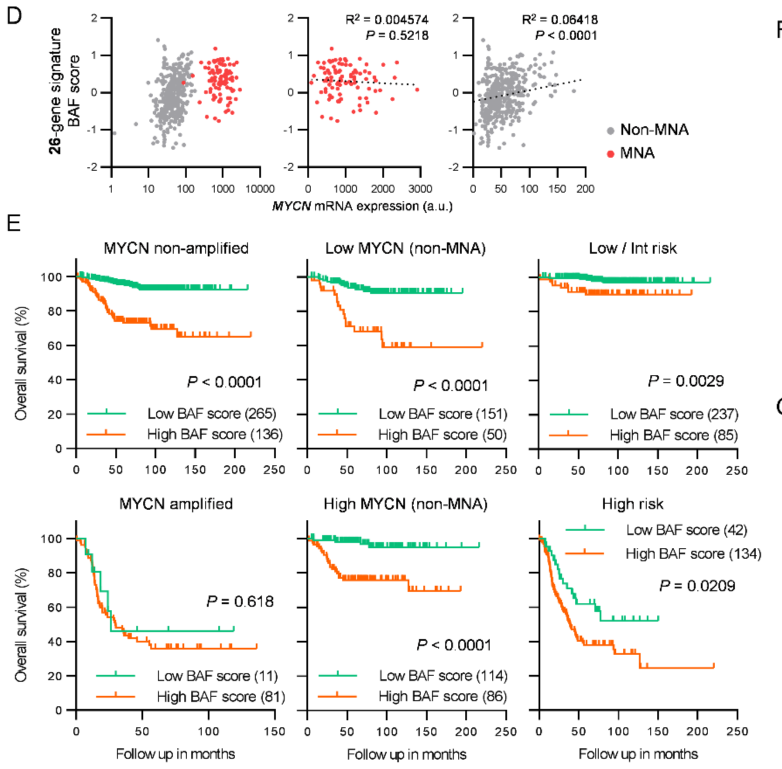


F

Univariate regression analysis (GSE62564)

| Factors | Overall survival | |
|---|---|---|
| | HR (95% CI) | *p* |
| Sex (M vs F) | - | 0.252 |
| Age (≥18 vs <18 months) | 8.114 (4.980-13.221) | 0.000 |
| MYCN (Amp vs no amp) | 7.793 (5.262-11.542) | 0.000 |
| INSS Stage (4 vs others) | 8.660 (5.441-13.783) | 0.000 |
| BAF score (High vs low) | 6.630 (4.105-10.708) | 0.000 |
| Risk (High vs others) | 21.423 (11.932-38.464) | 0.000 |

G

Multivariate regression analysis (GSE62564)

| Factors | Overall survival | |
|---|---|---|
| *Model 1* | HR (95% CI) | *p* |
| Age (≥18 vs < 18 months) | 3.448 (2.004-5.930) | 0.000 |
| MYCN (Amp vs no amp) | 2.759 (1.821-4.178) | 0.000 |
| INSS Stage (4 vs others) | 2.684 (1.578-4.563) | 0.000 |
| BAF score (High vs low) | 2.792 (1.655-4.709) | 0.000 |
| *Model 2* | | |
| BAF score (High vs low) | 2.271 (1.365-3.779) | 0.002 |
| Risk (High vs others) | 14.971 (8.046-27.858) | 0.000 |

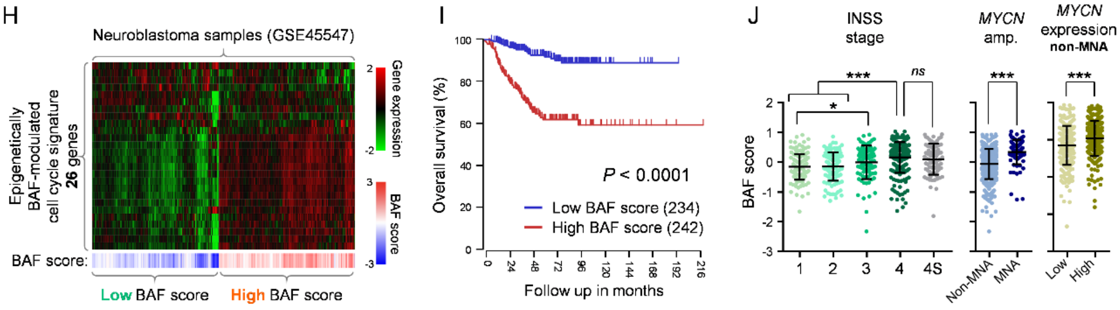

**Fig. S6**: Epigenetically BAF-modulated cell cycle signature keeps the prognostic predictive value in neuroblastoma. (**A**) mRNA expression of the BAF-modulated cell cycle transcriptional signature, narrowed down to those genes with associated ATAC-Seq chromatin remodeling repressive events (26 genes) in a public expression dataset of a 498 patient cohort (GSE62564). Patients were unbiasedly clustered into high and low epigenetic BAF score groups. (**B**) Kaplan-Meier plots comparing the overall survival of high and low epigenetic BAF score group of patients. (**C**) Comparison of epigenetic BAF signature score according to risk groups, INSS (International Neuroblastoma Staging System) stages, *MYCN* amplification (MNA), and *MYCN* mRNA expression (above or below median) in non-MNA cases. (**D**) Regression analyses between *MYCN* mRNA expression and epigenetic BAF score in GSE62564 dataset, separating groups according to *MYCN* amplification (MNA). (**E**) Kaplan-Meier plots comparing the overall survival between patients (GSE62564) with high or low epigenetic BAF score in separated groups, according to MYCN amplification (MNA), *MYCN* mRNA expression levels (above or below median) in non-MNA cases, and risk stratification. (**F**) Cox univariate regression analysis of overall survival with different clinic-pathological features. (**G**) Cox multivariate regression analysis of overall survival confirms epigenetic BAF cell cycle signature score as an independent prognostic marker in neuroblastoma. (**H**) An independent cohort of 649 patients (GSE45547) was used to validate the prognostic value of epigenetic BAF cell cycle signature. Heatmap shows RNA expression of 26 genes from BAF signature, with patients unbiasedly clustered into high and low BAF score groups. (**I**) Kaplan-Meier plots comparing the overall survival of high and low epigenetic BAF score group of patients from GSE45547. (**J**) Comparison of epigenetic BAF score between INSS stages, *MYCN* amplification (MNA), and *MYCN* mRNA expression levels (above or below median) in non-MNA cases (GSE45547). *ns* means 'non-significant'; * means $P < 0.05$; *** means $P < 0.001$.

Figure S7:

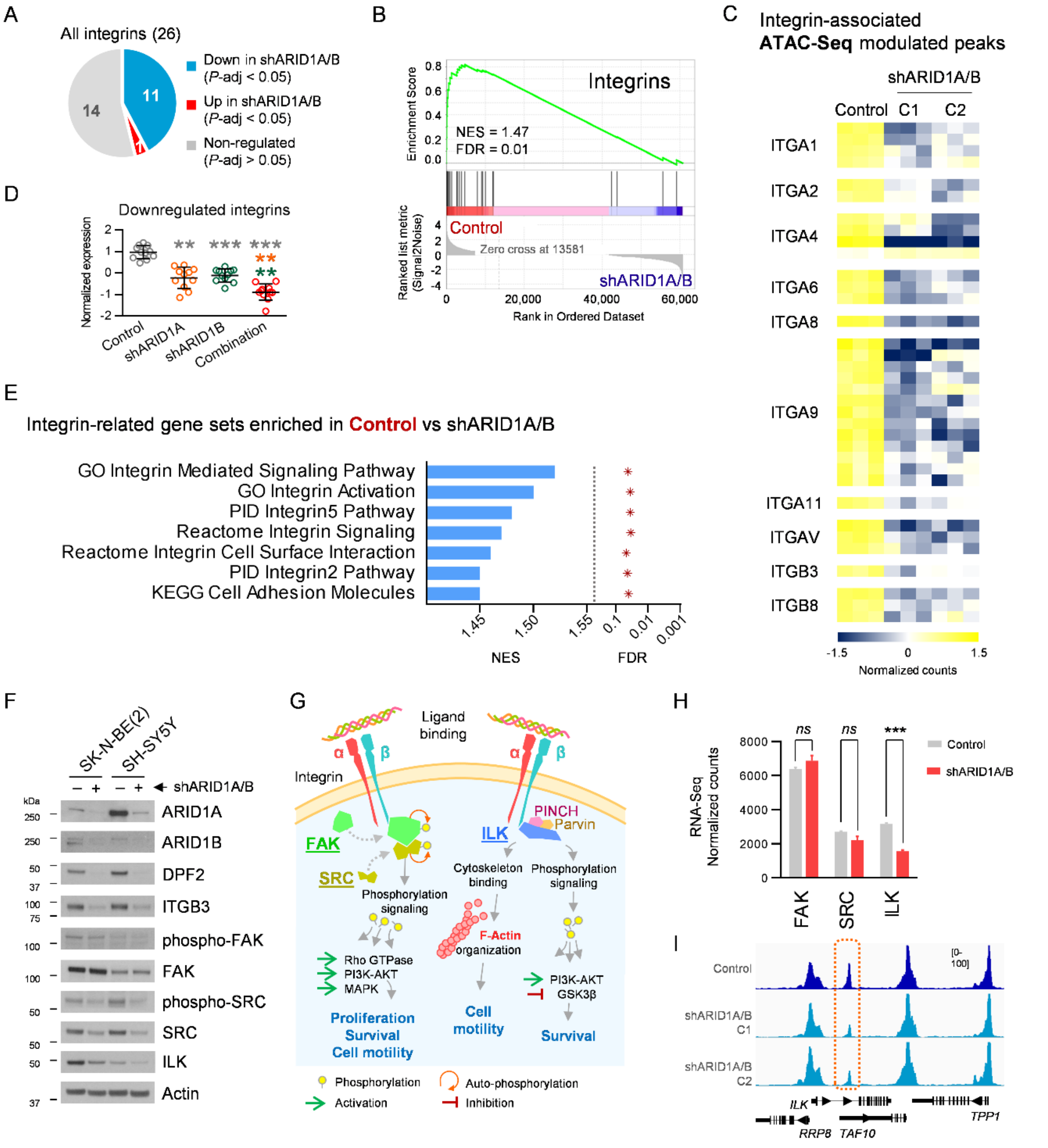


**Fig. S7**: BAF complex disruption promotes an epigenetic repression of the integrin gene family in neuroblastoma cells. (**A**) Pie chart of the proportion of human integrins down-, up- or non-regulated by BAF complex. (**B**) Enrichment plot of the 26 existing human integrins. NES and FDR are indicated. (**C**) Normalized peak coverage of the repressed ATAC-Seq peaks associated to 10 downregulated integrins after BAF disruption. (**D**) Normalized expression of the 11 repressed integrins after BAF inhibition after single or combined silencing of ARID1A and ARID1B. (**E**) Normalized Enrichment Score (NES) and False Discovery Rate (FDR) of the 5 integrin-related gene sets significantly enriched in RNA-Seq expression data of control compared to BAF-depleted SK-N-BE(2) cells. (**F**) Western blot analysis of different key elements involved in integrin signaling, 96 hours after BAF disruption. (**G**) Simplified scheme of the two main downstream integrin signaling pathways, adapted from [19] and [20]. (**H**) mRNA expression levels of *FAK*, *SRC* and *ILK* genes in BAF-disrupted SK-N-BE(2), from RNA-Seq data. (**I**) Representative genome browser ATAC-Seq coverage images of the repressed site associated to *ILK* gene. ns means 'non-significant'; *** means P < 0.001.

Figure S8:

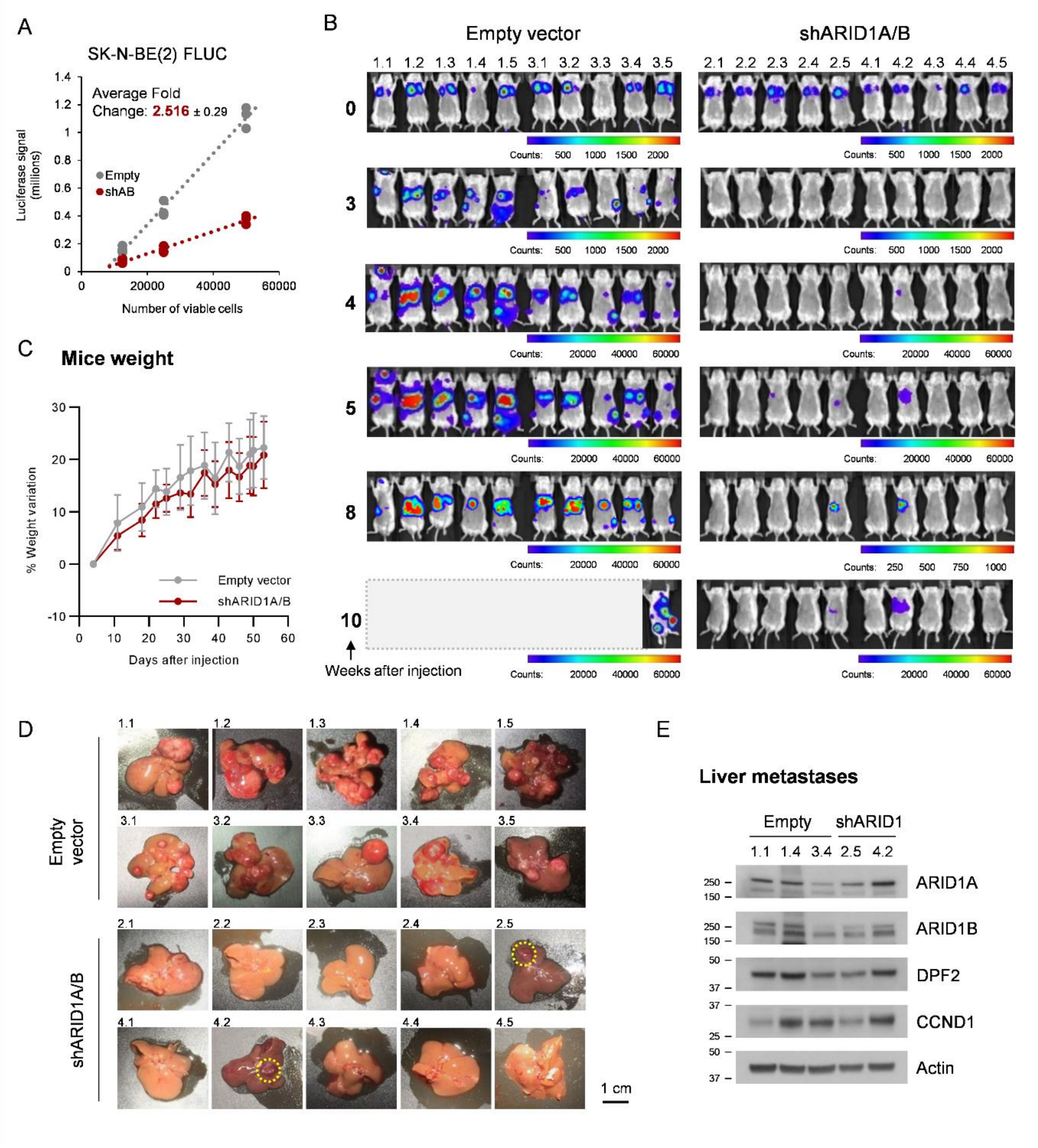


**Fig. S8**: Effects of BAF disruption on neuroblastoma macro-metastasis growth. (**A**) *In vitro* luciferase assay comparing luminescence signal of different numbers of pLKO-empty or pLKO-shARID1A/shARID1B (shARID1A/B or shAB) transduced SK-N-BE(2) FLUC viable cells. (**B**) Complete set of *in vivo* luminescence images of the long-term neuroblastoma metastasis model, at the indicated time points. Scale bar represents luminescence counts. (**C**) Mouse weight variation along the long-term metastasis experiment, starting from the day of cell injection. (**D**) Complete set of liver images at necropsy. Doted yellow circles denote macro-metastasis formed in the shARID1A/B group of mice. (**E**) Protein levels analysis of the indicated proteins of 3 empty-vector and 2 shARID1A/B liver macro-metastases, by western blot.

Figure S9:

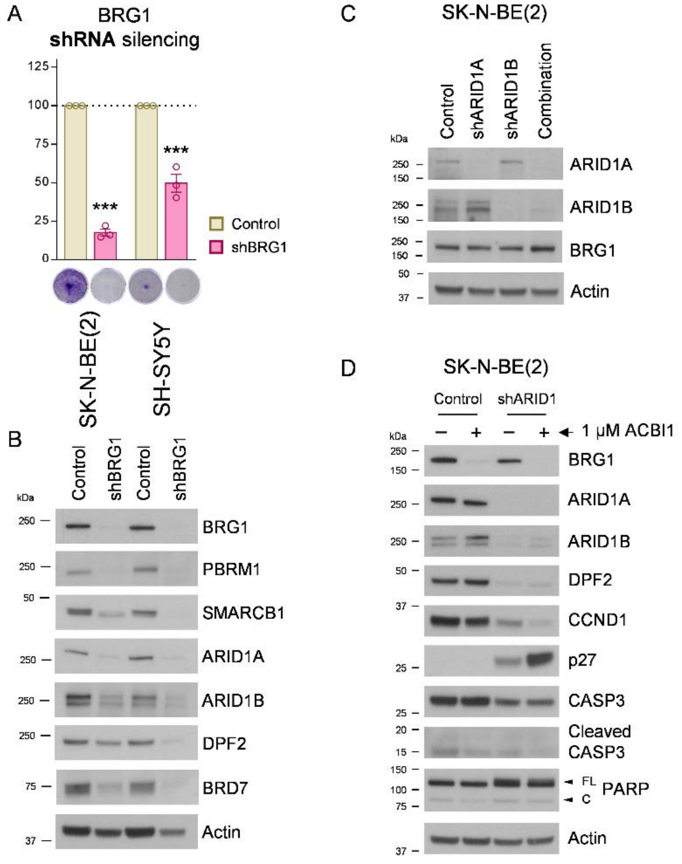


**Fig. S9**: Effects of shRNA-mediated BRG1 silencing in neuroblastoma cell lines. (**A**) Proliferation assays of SK-N-BE(2) and SH-SY5Y cells seeded 72 h post-transduction in 24-well plates and grown for 96 h. (**B**) Western blot analysis of BRG1 and other mSWI/SNF subunits 96 h after transduction with shBRG1 or control. (**C**) Western blot analysis of BRG1 protein levels after single or combined silencing of ARID1A and ARID1B (96 h after transduction) in SK-N-BE(2) cells. (**D**) Western blot analysis of apoptosis and cell cycle checkpoint regulators in SK-N-BE(2) transduced with shARID1A and shARID1B for 120 h, and treated with 1 µM ACBI1 (or control cis-ACBI1) for 96 h. FL means ‘Full Length’; C means ‘Cleaved’; *** means $P < 0.001$.

### 3. **Supplementary Tables**

**Table S1**: Mass Spectrometry-detected BRG1 interactors. Table shows those proteins significantly enriched in BRG1 (SMARCA4) immunoprecipitation in comparison with control IgG (BDFR < 0.05), in at least one of the two cell lines analysed (SK-N-BE(2) and SH-SY5Y). Coloured rows indicate interactors consistently identified in both cell lines.

| | | SK-N-BE(2) | | | | | | SH-SY5Y | | | | | |
|---|---|---|---|---|---|---|---|---|---|---|---|---|---|
| *Protein* | *UniProt ID* | *Spec* | *Spec Sum* | *Avg Spec* | *IgG Counts* | *Fold Change* | *BFDR* | *Spec* | *Spec Sum* | *Avg Spec* | *IgG Counts* | *Fold Change* | *BFDR* |
| **SMARCB1** | Q12824 | 30\|36\|33 | 99 | 33 | 0\|0\|0 | 330 | < 0.01 | 45\|44\|48 | 137 | 45.67 | 0\|1\|1 | 68.5 | < 0.01 |
| **SMARCC1** | Q92922 | 121\|126\|109 | 356 | 118.67 | 0\|1\|0 | 356 | < 0.01 | 136\|147\|152 | 435 | 145 | 1\|4\|3 | 54.38 | < 0.01 |
| **SMARCC2** | Q8TAQ2 | 107\|97\|97 | 301 | 100.33 | 0\|0\|0 | 1003.33 | < 0.01 | 122\|122\|131 | 375 | 125 | 1\|5\|5 | 34.09 | < 0.01 |
| **SMARCD1** | Q96GM5 | 63\|65\|63 | 191 | 63.67 | 0\|0\|0 | 636.67 | < 0.01 | 74\|74\|93 | 241 | 80.33 | 0\|0\|0 | 803.33 | < 0.01 |
| **SMARCD2** | Q92925 | 48\|48\|46 | 142 | 47.33 | 0\|0\|0 | 473.33 | < 0.01 | 52\|50\|52 | 154 | 51.33 | 0\|0\|0 | 513.33 | < 0.01 |
| **SMARCD3** | Q6STE5 | 39\|39\|40 | 118 | 39.33 | 0\|0\|0 | 393.33 | < 0.01 | 46\|45\|46 | 137 | 45.67 | 0\|0\|0 | 456.67 | < 0.01 |
| **SMARCE1** | Q969G3 | 64\|83\|58 | 205 | 68.33 | 5\|2\|2 | 22.78 | < 0.01 | 73\|74\|87 | 234 | 78 | 2\|4\|5 | 21.27 | < 0.01 |
| **ACTB** | P60709 | 59\|62\|61 | 182 | 60.67 | 11\|6\|11 | 6.5 | < 0.01 | 65\|77\|71 | 213 | 71 | 26\|13\|11 | 4.26 | < 0.01 |
| **ACTL6A** | O96019 | 45\|55\|57 | 157 | 52.33 | 0\|0\|0 | 523.33 | < 0.01 | 51\|57\|61 | 169 | 56.33 | 0\|0\|0 | 563.33 | < 0.01 |
| **ACTL6B** | O94805 | 4\|8\|10 | 22 | 7.33 | 0\|0\|0 | 73.33 | < 0.01 | 41\|47\|55 | 143 | 47.67 | 0\|0\|0 | 476.67 | < 0.01 |
| **BCL7A** | Q4VC05 | 21\|22\|20 | 63 | 21 | 0\|0\|0 | 210 | < 0.01 | 29\|26\|27 | 82 | 27.33 | 0\|0\|0 | 273.33 | < 0.01 |
| **BCL7B** | Q9BQE9 | 8\|7\|5 | 20 | 6.67 | 0\|0\|0 | 66.67 | < 0.01 | 8\|9\|7 | 24 | 8 | 0\|0\|0 | 80 | < 0.01 |
| **BCL7C** | Q8WUZ0 | 9\|6\|8 | 23 | 7.67 | 0\|0\|0 | 76.67 | < 0.01 | 7\|7\|8 | 22 | 7.33 | 0\|0\|0 | 73.33 | < 0.01 |
| **SMARCA4** | P51532 | 160\|151\|164 | 475 | 158.33 | 2\|2\|2 | 79.17 | < 0.01 | 182\|199\|196 | 577 | 192.33 | 2\|3\|3 | 72.13 | < 0.01 |
| **ARID1A** | O14497 | 145\|141\|145 | 431 | 143.67 | 0\|0\|0 | 1436.67 | < 0.01 | 196\|218\|219 | 633 | 211 | 0\|2\|1 | 211 | < 0.01 |
| **ARID1B** | Q8NFD5 | 103\|108\|93 | 304 | 101.33 | 0\|0\|0 | 1013.33 | < 0.01 | 113\|109\|116 | 338 | 112.67 | 0\|0\|0 | 1126.67 | < 0.01 |
| **DPF1** | Q92782 | 14\|10\|8 | 32 | 10.67 | 0\|0\|0 | 106.67 | < 0.01 | 21\|22\|26 | 69 | 23 | 0\|0\|0 | 230 | < 0.01 |
| **DPF2** | Q92785 | 42\|44\|39 | 125 | 41.67 | 0\|0\|0 | 416.67 | < 0.01 | 40\|45\|50 | 135 | 45 | 0\|2\|0 | 67.5 | < 0.01 |
| **DPF3** | Q92784 | 4\|4\|4 | 12 | 4 | 0\|0\|0 | 40 | < 0.01 | 20\|21\|19 | 60 | 20 | 0\|0\|0 | 200 | < 0.01 |
| **SS18** | Q15532 | 12\|12\|11 | 35 | 11.67 | 0\|0\|0 | 116.67 | < 0.01 | 12\|14\|12 | 38 | 12.67 | 0\|0\|0 | 126.67 | < 0.01 |
| **SS18L1** | O75177 | 10\|10\|10 | 30 | 10 | 0\|0\|0 | 100 | < 0.01 | 13\|13\|15 | 41 | 13.67 | 0\|0\|0 | 136.67 | < 0.01 |
| **ARID2** | Q68CP9 | 56\|57\|57 | 170 | 56.67 | 0\|1\|1 | 85 | < 0.01 | 48\|52\|64 | 164 | 54.67 | 3\|1\|0 | 41 | < 0.01 |
| **BRD7** | Q9NPI1 | 45\|50\|55 | 150 | 50 | 0\|0\|0 | 500 | < 0.01 | 40\|51\|58 | 149 | 49.67 | 0\|0\|0 | 496.67 | < 0.01 |
| **PBRM1** | Q86U86 | 95\|102\|92 | 289 | 96.33 | 0\|0\|0 | 963.33 | < 0.01 | 108\|118\|120 | 346 | 115.33 | 0\|0\|0 | 1153.33 | < 0.01 |
| **PHF10** | Q8WUB8 | 29\|26\|24 | 79 | 26.33 | 0\|0\|0 | 263.33 | < 0.01 | 28\|24\|29 | 81 | 27 | 0\|0\|0 | 270 | < 0.01 |
| **BICRA** | Q9NZM4 | 18\|18\|15 | 51 | 17 | 0\|0\|0 | 170 | < 0.01 | 24\|21\|25 | 70 | 23.33 | 0\|0\|0 | 233.33 | < 0.01 |
| **BICRAL** | Q6AI39 | 16\|14\|10 | 40 | 13.33 | 0\|0\|0 | 133.33 | < 0.01 | 17\|17\|17 | 51 | 17 | 0\|0\|0 | 170 | < 0.01 |
| **BRD9** | Q9H8M2 | 12\|7\|9 | 28 | 9.33 | 0\|0\|0 | 93.33 | < 0.01 | 14\|15\|24 | 53 | 17.67 | 0\|0\|0 | 176.67 | < 0.01 |
| **ACTA1** | P68133 | 41\|44\|36 | 121 | 40.33 | 10\|5\|9 | 5.04 | < 0.01 | 50\|55\|52 | 157 | 52.33 | 20\|12\|11 | 3.65 | < 0.01 |
| ACTBL2 | Q562R1 | | | Undetected | | | | 21\|22\|20 | 63 | 21 | 10\|6\|5 | 3 | < 0.01 |
| **ACTG1** | P63261 | 62\|64\|64 | 190 | 63.33 | 12\|7\|11 | 6.33 | < 0.01 | 66\|80\|72 | 218 | 72.67 | 26\|14\|12 | 4.19 | < 0.01 |
| CAPRIN1 | Q14444 | 1\|3\|4 | 8 | 2.67 | 0\|1\|0 | 8 | 0.02 | 1\|4\|6 | 11 | 3.67 | 0\|0\|2 | 5.5 | 0.1 |
| DDX21 | Q9NR30 | 6\|4\|8 | 18 | 6 | 10\|1\|2 | 1.38 | 0.65 | 6\|9\|10 | 25 | 8.33 | 2\|2\|1 | 5 | < 0.01 |
| FBL | P22087 | 0\|1\|1 | 2 | 0.67 | 0\|0\|0 | 6.67 | 0.74 | 2\|2\|2 | 6 | 2 | 0\|1\|0 | 6 | 0.02 |
| G3BP2 | Q9UN86 | 1\|3\|2 | 6 | 2 | 0\|0\|0 | 20 | 0.03 | 2\|1\|1 | 4 | 1.33 | 0\|0\|0 | 13.33 | 0.25 |
| **HSPA5** | P11021 | 9\|7\|7 | 23 | 7.67 | 0\|0\|0 | 76.67 | < 0.01 | 11\|11\|10 | 32 | 10.67 | 0\|0\|0 | 106.67 | < 0.01 |
| KHDRBS1 | Q07666 | 2\|1\|3 | 6 | 2 | 2\|1\|0 | 2 | 0.44 | 6\|6\|7 | 19 | 6.33 | 1\|3\|0 | 4.75 | 0.01 |
| KHDRBS2 | Q5VWX1 | | | Undetected | | | | 2\|3\|2 | 7 | 2.33 | 0\|1\|0 | 7 | 0.02 |
| **KPNA2** | P52292 | 10\|10\|10 | 30 | 10 | 0\|1\|1 | 15 | < 0.01 | 16\|14\|17 | 47 | 15.67 | 2\|1\|0 | 15.67 | < 0.01 |
| NCL | P19338 | 25\|24\|33 | 82 | 27.33 | 26\|7\|16 | 1.67 | 0.71 | 24\|24\|30 | 78 | 26 | 9\|7\|8 | 3.25 | < 0.01 |
| PABPC4 | Q13310 | | | Undetected | | | | 2\|2\|5 | 9 | 3 | 0\|1\|0 | 9 | 0.02 |
| **PCBP2** | Q15366 | 3\|5\|3 | 11 | 3.67 | 2\|0\|0 | 5.5 | < 0.01 | 5\|5\|6 | 16 | 5.33 | 2\|2\|0 | 4 | 0.01 |
| PSIP1 | O75475 | | | Undetected | | | | 4\|3\|4 | 11 | 3.67 | 1\|2\|0 | 3.67 | 0.04 |
| RBM15 | Q96T37 | 4\|2\|1 | 7 | 2.33 | 1\|1\|2 | 1.75 | 0.44 | 5\|5\|6 | 16 | 5.33 | 2\|2\|0 | 4 | 0.01 |
| **RPL10A** | P62906 | 4\|0\|4 | 8 | 2.67 | 1\|0\|0 | 8 | 0.01 | 3\|5\|5 | 13 | 4.33 | 0\|1\|1 | 6.5 | 0.01 |
| RPL18A | Q02543 | 3\|3\|3 | 9 | 3 | 2\|3\|1 | 1.5 | 0.6 | 3\|4\|4 | 11 | 3.67 | 2\|0\|1 | 3.67 | 0.04 |
| RPL6 | Q02878 | 7\|9\|6 | 22 | 7.33 | 4\|0\|4 | 2.75 | 0.21 | 10\|8\|7 | 25 | 8.33 | 2\|1\|2 | 5 | < 0.01 |
| RPLP2 | P05387 | 5\|3\|2 | 10 | 3.33 | 1\|0\|2 | 3.33 | 0.11 | 4\|6\|5 | 15 | 5 | 0\|0\|0 | 50 | < 0.01 |
| RPS12 | P25398 | 4\|2\|5 | 11 | 3.67 | 0\|0\|0 | 36.67 | < 0.01 | 0\|1\|5 | 6 | 2 | 0\|0\|0 | 20 | 0.22 |
| RPS20 | P60866 | 3\|3\|3 | 9 | 3 | 1\|1\|1 | 3 | 0.17 | 4\|4\|3 | 11 | 3.67 | 0\|1\|0 | 11 | < 0.01 |
| RPS24 | P62847 | 1\|1\|2 | 4 | 1.33 | 2\|2\|2 | 0.67 | 0.74 | 3\|4\|2 | 9 | 3 | 1\|1\|0 | 4.5 | 0.03 |
| SRP72 | O76094 | 0\|1\|1 | 2 | 0.67 | 0\|0\|1 | 2 | 0.74 | 2\|4\|3 | 9 | 3 | 1\|0\|1 | 4.5 | 0.03 |
| SRPRA | P08240 | 0\|2\|2 | 4 | 1.33 | 0\|0\|0 | 13.33 | 0.04 | 1\|1\|0 | 2 | 0.67 | 0\|0\|0 | 6.67 | 0.71 |
| SYNCRIP | O60506 | 1\|3\|7 | 11 | 3.67 | 5\|1\|1 | 1.57 | 0.45 | 7\|8\|8 | 23 | 7.67 | 1\|3\|2 | 3.83 | < 0.01 |
| U2AF1L5 | P0DN76 | 2\|0\|0 | 2 | 0.67 | 2\|0\|0 | 1 | 0.46 | 3\|4\|4 | 11 | 3.67 | 0\|1\|0 | 11 | < 0.01 |

Spec: Spectral counts; Avg Spec: Average Spectral counts; BFDR: Bayesian False Discovery Rate.

**Table S2**: Gene Set Enrichment Analysis of 50 Hallmarks after BAF disruption. Hallmark collection was extracted from MSigDB, and comparison of RNA-Seq data from shARID1A/B (Combination) against Control cells was performed.

| *Gene Set* | *Size* | *ES* | *NES* | *NOM p-val* | *FDR q-val* |
|---|---|---|---|---|---|
| HALLMARK_EPITHELIAL_MESENCHYMAL_TRANSITION | 200 | 0.71 | 1.53 | 0 | 0.014 |
| HALLMARK_E2F_TARGETS | 200 | 0.86 | 1.52 | 0 | 0.014 |
| HALLMARK_INTERFERON_ALPHA_RESPONSE | 97 | 0.6 | 1.51 | 0 | 0.014 |
| HALLMARK_G2M_CHECKPOINT | 200 | 0.82 | 1.49 | 0 | 0.014 |
| HALLMARK_MITOTIC_SPINDLE | 199 | 0.69 | 1.48 | 0 | 0.016 |
| HALLMARK_MYOGENESIS | 200 | 0.49 | 1.46 | 0 | 0.016 |
| HALLMARK_ANDROGEN_RESPONSE | 100 | 0.62 | 1.46 | 0 | 0.016 |
| HALLMARK_APOPTOSIS | 161 | 0.55 | 1.45 | 0 | 0.018 |
| HALLMARK_INTERFERON_GAMMA_RESPONSE | 200 | 0.51 | 1.43 | 0 | 0.019 |
| HALLMARK_NOTCH_SIGNALING | 32 | 0.59 | 1.41 | 0 | 0.019 |
| HALLMARK_INFLAMMATORY_RESPONSE | 200 | 0.51 | 1.44 | 0 | 0.02 |
| HALLMARK_KRAS_SIGNALING_UP | 200 | 0.55 | 1.43 | 0 | 0.02 |
| HALLMARK_IL2_STAT5_SIGNALING | 199 | 0.51 | 1.4 | 0 | 0.02 |
| HALLMARK_APICAL_JUNCTION | 200 | 0.53 | 1.39 | 0 | 0.024 |
| HALLMARK_GLYCOLYSIS | 200 | 0.56 | 1.38 | 0 | 0.04 |
| HALLMARK_HYPOXIA | 200 | 0.55 | 1.37 | 0 | 0.043 |
| HALLMARK_UV_RESPONSE_DN | 144 | 0.54 | 1.36 | 0 | 0.044 |
| HALLMARK_CHOLESTEROL_HOMEOSTASIS | 74 | 0.55 | 1.36 | 0 | 0.046 |
| HALLMARK_COMPLEMENT | 200 | 0.48 | 1.34 | 0 | 0.05 |
| HALLMARK_COAGULATION | 138 | 0.48 | 1.33 | 0 | 0.052 |
| HALLMARK_ESTROGEN_RESPONSE_LATE | 200 | 0.53 | 1.33 | 0 | 0.059 |
| HALLMARK_TGF_BETA_SIGNALING | 54 | 0.58 | 1.32 | 0.024 | 0.064 |
| HALLMARK_ANGIOGENESIS | 36 | 0.54 | 1.31 | 0 | 0.071 |
| HALLMARK_TNFA_SIGNALING_VIA_NFKB | 200 | 0.52 | 1.29 | 0.05 | 0.082 |
| HALLMARK_IL6_JAK_STAT3_SIGNALING | 87 | 0.48 | 1.28 | 0.026 | 0.082 |
| HALLMARK_APICAL_SURFACE | 44 | 0.51 | 1.29 | 0 | 0.084 |
| HALLMARK_PROTEIN_SECRETION | 96 | 0.52 | 1.28 | 0.054 | 0.085 |
| HALLMARK_SPERMATOGENESIS | 135 | 0.43 | 1.27 | 0.079 | 0.088 |
| HALLMARK_DNA_REPAIR | 149 | 0.51 | 1.27 | 0.028 | 0.092 |
| HALLMARK_WNT_BETA_CATENIN_SIGNALING | 42 | 0.48 | 1.25 | 0.038 | 0.102 |
| HALLMARK_HEME_METABOLISM | 200 | 0.4 | 1.21 | 0 | 0.124 |
| HALLMARK_PI3K_AKT_MTOR_SIGNALING | 105 | 0.46 | 1.22 | 0.125 | 0.125 |
| HALLMARK_UV_RESPONSE_UP | 157 | 0.49 | 1.21 | 0 | 0.126 |
| HALLMARK_ESTROGEN_RESPONSE_EARLY | 200 | 0.45 | 1.22 | 0.029 | 0.127 |
| HALLMARK_ADIPOGENESIS | 199 | 0.39 | 1.19 | 0 | 0.148 |
| HALLMARK_MYC_TARGETS_V1 | 200 | 0.58 | 1.17 | 0.382 | 0.162 |
| HALLMARK_ALLOGRAFT_REJECTION | 200 | 0.38 | 1.15 | 0 | 0.2 |
| HALLMARK_XENOBIOTIC_METABOLISM | 200 | 0.38 | 1.14 | 0.092 | 0.21 |
| HALLMARK_HEDGEHOG_SIGNALING | 36 | 0.46 | 1.09 | 0.286 | 0.255 |
| HALLMARK_FATTY_ACID_METABOLISM | 158 | 0.33 | 1.09 | 0.114 | 0.255 |
| HALLMARK_P53_PATHWAY | 200 | 0.37 | 1.07 | 0.28 | 0.277 |
| HALLMARK_MTORC1_SIGNALING | 200 | 0.46 | 0.99 | 0.534 | 0.441 |
| HALLMARK_PANCREAS_BETA_CELLS | 40 | 0.37 | 0.98 | 0.523 | 0.45 |
| HALLMARK_MYC_TARGETS_V2 | 58 | 0.52 | 0.97 | 0.538 | 0.466 |
| HALLMARK_UNFOLDED_PROTEIN_RESPONSE | 113 | 0.46 | 0.97 | 0.646 | 0.471 |
| HALLMARK_BILE_ACID_METABOLISM | 112 | 0.33 | 0.86 | 0.79 | 0.702 |
| HALLMARK_REACTIVE_OXYGEN_SPECIES_PATHWAY | 49 | 0.37 | 0.78 | 0.828 | 0.832 |
| HALLMARK_PEROXISOME | 104 | -0.35 | -1.13 | 0.044 | 0.565 |
| HALLMARK_KRAS_SIGNALING_DN | 200 | -0.31 | -0.93 | 0.721 | 0.836 |
| HALLMARK_OXIDATIVE_PHOSPHORYLATION | 200 | -0.38 | -0.88 | 0.649 | 0.663 |

ES: Enrichment Score; NES: Normalized Enrichment Score; NOM *p*-val: Nominal *p* value; FDR *q*-val: False Discovery Rate *q*-value.

**Table S3**: BAF-modulated cell cycle gene signature (171 genes).

| # | *Gene Symbol* | # | *Gene Symbol* | # | *Gene Symbol* |
|---|---|---|---|---|---|
| 1 | *EZR* | 58 | *BUB3* | 115 | *CDC25A* |
| 2 | *NEDD9* | 59 | *NET1* | 116 | *NDC80* |
| 3 | *MYC* | 60 | *RBM14* | 117 | *TTK* |
| 4 | *MYO9B* | 61 | *LBR* | 118 | *KIF23* |
| 5 | *PCGF5* | 62 | *DEK* | 119 | *BUB1* |
| 6 | *PALLD* | 63 | *SYNCRIP* | 120 | *CENPE* |
| 7 | *ACTN4* | 64 | *MMS22L* | 121 | *PBK* |
| 8 | *VCL* | 65 | *CASP8AP2* | 122 | *ECT2* |
| 9 | *MYH9* | 66 | *STAG1* | 123 | *CDK1* |
| 10 | *FLNA* | 67 | *SMC3* | 124 | *CCNA2* |
| 11 | *MYO1E* | 68 | *MSH2* | 125 | *ANLN* |
| 12 | *PREX1* | 69 | *G3BP1* | 126 | *INCENP* |
| 13 | *NOTCH2* | 70 | *CDC7* | 127 | *WEE1* |
| 14 | *EPB41L2* | 71 | *CKS1B* | 128 | *KNTC1* |
| 15 | *DOCK2* | 72 | *DUT* | 129 | *ESPL1* |
| 16 | *SLC12A2* | 73 | *SMC1A* | 130 | *TROAP* |
| 17 | *FLNB* | 74 | *KPNB1* | 131 | *KIF18B* |
| 18 | *SMAD3* | 75 | *UBR7* | 132 | *MCM4* |
| 19 | *WDR90* | 76 | *CDKN2C* | 133 | *POLA2* |
| 20 | *PKD2* | 77 | *USP1* | 134 | *TK1* |
| 21 | *SHROOM1* | 78 | *TCF19* | 135 | *BIRC5* |
| 22 | *CUL1* | 79 | *E2F2* | 136 | *RRM2* |
| 23 | *EFNA5* | 80 | *KIF2C* | 137 | *LMNB1* |
| 24 | *CCND1* | 81 | *CDCA8* | 138 | *TOP2A* |
| 25 | *STMN1* | 82 | *CIT* | 139 | *TMPO* |
| 26 | *TUBB* | 83 | *RFC2* | 140 | *KIF22* |
| 27 | *KIF3B* | 84 | *MCM7* | 141 | *MCM5* |
| 28 | *CLIP2* | 85 | *NUP107* | 142 | *CHAF1A* |
| 29 | *MYH10* | 86 | *SMC2* | 143 | *POLD1* |
| 30 | *RPA1* | 87 | *CDK5RAP2* | 144 | *NCAPD2* |
| 31 | *CDC42BPA* | 88 | *CENPJ* | 145 | *TIMELESS* |
| 32 | *DOCK4* | 89 | *NASP* | 146 | *MCM6* |
| 33 | *DST* | 90 | *EXO1* | 147 | *MCM2* |
| 34 | *CCDC88A* | 91 | *E2F3* | 148 | *MCM3* |
| 35 | *TRIO* | 92 | *KPNA2* | 149 | *TACC3* |
| 36 | *SSH2* | 93 | *CBX5* | 150 | *PCNA* |
| 37 | *DR1* | 94 | *CCNF* | 151 | *E2F1* |
| 38 | *SRSF10* | 95 | *ANP32E* | 152 | *MYBL2* |
| 39 | *MTF2* | 96 | *AURKB* | 153 | *HMGB2* |
| 40 | *XPO1* | 97 | *GINS2* | 154 | *PRC1* |
| 41 | *KIF5B* | 98 | *DDX39A* | 155 | *RACGAP1* |
| 42 | *PDS5B* | 99 | *PLK1* | 156 | *CENPF* |
| 43 | *RICTOR* | 100 | *UBE2C* | 157 | *TPX2* |
| 44 | *CNTRL* | 101 | *CCNB2* | 158 | *MKI67* |
| 45 | *NBN* | 102 | *UBE2T* | 159 | *NUSAP1* |
| 46 | *ARFGEF1* | 103 | *UBE2S* | 160 | *KIF11* |
| 47 | *NIN* | 104 | *RNASEH2A* | 161 | *BUB1B* |
| 48 | *ARF6* | 105 | *RANBP1* | 162 | *ATAD2* |
| 49 | *ARHGAP5* | 106 | *HMGB3* | 163 | *SMC4* |
| 50 | *FARP1* | 107 | *UNG* | 164 | *DNMT1* |
| 51 | *SLC38A1* | 108 | *TRA2B* | 165 | *SRSF2* |
| 52 | *NUDT21* | 109 | *HNRNPD* | 166 | *SRSF1* |
| 53 | *CCP110* | 110 | *BRCA1* | 167 | *EZH2* |
| 54 | *RFC1* | 111 | *CDC6* | 168 | *SPAG5* |
| 55 | *DCK* | 112 | *RFC3* | 169 | *POLE* |
| 56 | *SAC3D1* | 113 | *MAD2L1* | 170 | *LIG1* |
| 57 | *MXD3* | 114 | *KIF15* | 171 | *BRCA2* |

**Table S4**: BAF signature score (171 genes) correlations with clinical variables in neuroblastoma, using Fisher's test (GSE62564, n = 498).

| ***Variable*** | ***Σ*** | ***BAF score*** | | ***p value*** |
|---|---|---|---|---|
| | | ***Low*** | ***High*** | |
| All patients | 498 | 300 (60.2%) | 198 (39.8%) | |
| Sex | | | | |
| *M* | 287 | 170 (59.2%) | 117 (40.8%) | 0.643 |
| *F* | 211 | 130 (61.6%) | 81 (38.4%) | |
| Age | | | | |
| *<18 months* | 300 | 213 (71.0%) | 87 (29.0%) | 0.000 |
| *≥18 months* | 198 | 87 (43.9%) | 111 (56.1%) | |
| INSS Stage | | | | |
| *1* | 121 | 107 (88.4%) | 14 (11.6%) | 0.000 |
| *2* | 78 | 61 (78.2%) | 17 (21.8%) | |
| *3* | 63 | 38 (60.3%) | 25 (39.7%) | |
| *4* | 183 | 57 (31.1%) | 126 (68.9%) | |
| *4s* | 53 | 37 (69.8%) | 16 (30.2%) | |
| ISSN Stage | | | | |
| *1,2,3 & 4s* | 315 | 243 (77.1%) | 72 (22.9%) | 0.000 |
| *4* | 183 | 57 (31.1%) | 126 (68.9%) | |
| MYCN | | | | |
| *No amp* | 401 | 285 (71.1%) | 116 (28.9%) | 0.000 |
| *Amp* | 92 | 12 (13.0%) | 80 (87.0%) | |
| Risk | | | | |
| *Low, Intermediate* | 322 | 257 (79.8%) | 65 (20.2%) | 0.000 |
| *High* | 176 | 43 (24.4%) | 133 (75.6%) | |
| OS (Death) | | | | |
| *No* | 393 | 277 (70.5%) | 116 (29.5%) | 0.000 |
| *Yes* | 105 | 23 (21.9%) | 82 (78.1%) | |
| EFS (Recurrence) | | | | |
| *No* | 315 | 233 (74.0%) | 82 (26.0%) | 0.000 |
| *Yes* | 183 | 67 (36.6%) | 116 (63.4%) | |
| FAV | | | | |
| *Favourable* | 181 | 162 (89.5%) | 19 (10.5%) | 0.000 |
| *Unfavourable* | 91 | 22 (24.2%) | 69 (75.8%) | |

EFS: Event-free survival; FAV: Unfavourable/Favourable (class label for extreme disease course); HR: High-risk patients; OS: Overall survival.

**Table S5**: Epigenetically BAF-modulated cell cycle signature (26 genes).

| # | *Gene Symbol* | # | *Gene Symbol* | # | *Gene Symbol* |
|---|---|---|---|---|---|
| 1 | *FARP1* | 11 | *LMNB1* | 21 | *ARFGEF1* |
| 2 | *LBR* | 12 | *MKI67* | 22 | *CCDC88A* |
| 3 | *CDK5RAP2* | 13 | *TIMELESS* | 23 | *DOCK2* |
| 4 | *DEK* | 14 | *MCM3* | 24 | *NOTCH2* |
| 5 | *USP1* | 15 | *HMGB2* | 25 | *NEDD9* |
| 6 | *CDC25A* | 16 | *RRM2* | 26 | *SLC12A2* |
| 7 | *ANLN* | 17 | *CDC7* | | |
| 8 | *ECT2* | 18 | *KIF3B* | | |
| 9 | *RFC3* | 19 | *TRIO* | | |
| 10 | *MCM5* | 20 | *MYH10* | | |

**Table S6**: Epigenetic BAF signature score (26 genes) correlations with clinical variables in neuroblastoma, using Fisher's test (GSE62564, n = 498).

| ***Variable*** | **Σ** | ***Epigenetic BAF score*** | | ***p value*** |
|---|---|---|---|---|
| | | ***Low*** | ***High*** | |
| All patients | 498 | 279 (56.0%) | 219 (44.4%) | |
| Sex | | | | |
| *M* | 287 | 158 (51.1%) | 129 (44.9%) | 0.648 |
| *F* | 211 | 121 (57.3%) | 90 (42.7%) | |
| Age | | | | |
| *<18 months* | 300 | 194 (64.7%) | 106 (35.3%) | 0.000 |
| *≥18 months* | 198 | 85 (42.9%) | 113 (57.1%) | |
| INSS Stage | | | | |
| *1* | 121 | 102 (84.3%) | 19 (15.7%) | 0.000 |
| *2* | 78 | 57 (73.1%) | 21 (26.9%) | |
| *3* | 63 | 31 (49.2%) | 32 (50.8%) | |
| *4* | 183 | 55 (30.1%) | 128 (69.9%) | |
| *4s* | 53 | 34 (64.2%) | 19 (35.8%) | |
| ISSN Stage | | | | |
| *1,2,3 & 4s* | 315 | 224 (71.1%) | 91 (28.9%) | 0.000 |
| *4* | 183 | 55 (30.1%) | 128 (69.9%) | |
| MYCN | | | | |
| *No amp* | 401 | 265 (66.1%) | 136 (33.9%) | 0.000 |
| *Amp* | 92 | 11 (12.0%) | 81 (88.0%) | |
| Risk | | | | |
| *Low, Intermediate* | 322 | 237 (73.6%) | 85 (26.4%) | 0.000 |
| *High* | 176 | 42 (23.9%) | 134 (76.1%) | |
| OS (Death) | | | | |
| *No* | 393 | 258 (65.6%) | 135 (34.4%) | 0.000 |
| *Yes* | 105 | 21 (20.0%) | 84 (80.0%) | |
| EFS (Recurrence) | | | | |
| *No* | 315 | 217 (68.9%) | 98 (31.1%) | 0.000 |
| *Yes* | 183 | 62 (33.9%) | 121 (66.1%) | |
| FAV | | | | |
| *Favourable* | 181 | 152 (84.0%) | 29 (16.0%) | 0.000 |
| *Unfavourable* | 91 | 20 (22.0%) | 71 (78.0%) | |

EFS: Event-free survival; FAV: Unfavourable/Favourable (class label for extreme disease course); HR: High-risk patients; OS: Overall survival.

**Table S7**: Gene Ontology (Biological Process) categories enriched (Benjamini-corrected *p*-value < 0.05) analysis results on 469 ATAC and RNA-Seq repressed genes after BAF disruption.

| *Gene Ontology term* | *Number of genes* | p *value* |
|---|---|---|
| Extracellular matrix organization | 36 | $1.7 \times 10^{-14}$ |
| Cell adhesion | 40 | $1.8 \times 10^{-7}$ |
| Cell-matrix adhesion | 16 | $9.7 \times 10^{-6}$ |
| Cell adhesion mediated by integrin | 10 | $6.7 \times 10^{-5}$ |
| Angiogenesis | 22 | $7.4 \times 10^{-5}$ |
| Collagen fibril organization | 14 | $2.3 \times 10^{-4}$ |
| Integrin-mediated signaling pathway | 13 | $1.7 \times 10^{-3}$ |
| Cell-cell adhesion | 17 | $1.7 \times 10^{-3}$ |
| Positive regulation of angiogenesis | 15 | $3.9 \times 10^{-3}$ |
| Lung alveolus development | 8 | $5.2 \times 10^{-3}$ |
| Skeletal system development | 13 | $9.8 \times 10^{-3}$ |
| Modulation of synaptic transmission | 9 | 0.02 |
| Odontogenesis | 7 | 0.02 |
| Positive regulation of neuron projection development | 12 | 0.02 |
| Cell morphogenesis | 10 | 0.02 |
| Positive regulation of cell-substrate adhesion | 7 | 0.03 |
| Cell-cell junction assembly | 7 | 0.03 |
| Substrate-dependent cell migration | 4 | 0.03 |
| Response to hypoxia | 14 | 0.03 |
| Positive regulation of apoptotic process | 21 | 0.04 |
| Wound healing | 10 | 0.04 |

**Table S8**: Epigenetically repressed genes after BAF disruption included in the enriched Gene Ontology (GO) categories (Benjamini corrected *p*-value < 0.01).

| *GO term* | *Enriched genes* |
|---|---|
| **ECM organization** | *ABI3BP, ADAM12, ADAMTS14, ADAMTS15, ADAMTS2, ADAMTS9, SH3PXD2A, CCDC80, COLQ, COL1A2, COL2A1, COL4A1, COL9A3, COL5A1, COL6A3, COL22A1, DMP1, FBLN2, ITGA11, ITGA2, ITGA4, ITGA6, ITGA8, ITGA9, ITGAV, ITGB3, ITGB6, LUM, MMP14, NID1, OLFML2A, OLFML2B, PTX3, TGFBI, VCAM1, VCAN* |
| **Cell adhesion** | *ADAM12, CD177, CD226, CD34, CD9, CD96, EPHA4, PDZD2, CDH11, CDH2, CDH6, COL5A1, COL6A3, DPT, EMP2, ISLR, IGFBP7, ITGA11, ITGA2, ITGA4, ITGA6, ITGA8, ITGA9, ITGAV, ITGB3, ITGB6, LOXL2, MAGI1, MYH10, NEDD9, NTM, OPCML, PCDH7, RELN, SEMA5A, SRPX, TGFBI, TPBG, VCAM1, VCAN* |
| **Cell-matrix adhesion** | *CD34, CD96, FREM1, EMP2, ILK, ITGA11, ITGA2, ITGA4, ITGA6, ITGA8, ITGA9, ITGAV, ITGB3, ITGB6, NID1, VCAM1* |
| **Cell adhesion by integrin** | *EXT1, ITGA11, ITGA2, ITGA4, ITGA6, ITGA8, ITGA9, ITGAV, ITGB3, ITGB6* |
| **Angiogenesis** | *GAB1, RAPGEF3, ANGPT1, ANGPT2, ANGPTL4, APLNR, CALCRL, CSPG4, COL22A1, EMCN, EDN2, FGFR2, GJA5, HS6ST1, ITGAV, MMP14, PDGFRB, RBPJ, THSD7A, TGFBI, TMEM100, VASH1* |
| **Collagen organization** | *ADAMTS14, ADAMTS2, COL1A2, COL2A1, COL4A1, COL9A3, COL5A1, COL6A3, COL22A1, DPT, EXT1, ITGA6, LUM, LOXL2* |
| **Integrin signaling** | *ADAM12, ILK, ITGA11, ITGA2, ITGA4, ITGA6, ITGA8, ITGA9, ITGAV, ITGB3, ITGB6, NEDD9, PTN* |
| **Cell-cell adhesion** | *BCL2, CD34, LPP, CDH11, CDH2, CDH5, EMCN, ITGA11, ITGA2, ITGA4, ITGA6, ITGA8, ITGA9, ITGAV, NEGR1, TJP2, VCAM1* |
| **Positive angiogenesis** | *ADAM12, CD34, ETS1, GAB1, RAPGEF3, ANGPT2, ANGPTL4, APLNR, AQP1, CDH5, HGF, NTRK1, SEMA5A, TWIST1, VEGFC* |
| **Lung alveolus development** | *ERRFI1, BMP4, EDN2, FGFR2, HS6ST1, ITGB6, MAN1A2, MAN2A1* |
| **Skeletal development** | *ETS2, CDH11, CMKLR1, COL1A2, COL2A1, EXT1, FGFR1, FRZB, MMP14, RPS6KA3, USP1, VCAN, ZBTB16* |

**Table S9**: shRNA and siRNA sequences.

| *Target* | *Name* | *Target sequence* | *Transcript* | *Source* |
|---|---|---|---|---|
| ***shRNA*** | | | | |
| ARID1A | shARID1A #2 | GCCTGATCTATCTGGTTCAAT | NM_006015.6 | Mission shRNA TRCN0000059089 |
| | shARID1A #4 | CCGTTGATGAACTCATTGGTT | | Mission shRNA TRCN0000059091 |
| ARID1B | shARDI1B #2 | GGGTTTGGCCCAGGTTAATAA | NM_017519.3 | Mission shRNA TRCN0000416443 |
| | shARID1B #3 | TGCTGTCTAGTGCATTCAAAG | | Mission shRNA TRCN0000436265 |
| ARID2 | shARID2 #3 | CGTACCTGTCTTCGTTTCCTA | NM_152641.4 | Mission shRNA TRCN0000166264 |
| | shARID2 #4 | CCTCCTTCAAACTCAGGGAAA | | Mission shRNA TRCN0000166321 |
| SMARCC1 | shSMARCC1 #2 | GCTATGATACTTGGGTCCATA | NM_003074.4 | Mission shRNA TRCN0000278033 |
| | shSMARCC1 #5 | GCTATGATACTTGGGTCCATA | | Mission shRNA TRCN0000015630 |
| SMARCC2 | shSMARCC2 #2 | TCACTAAACTGCCGATCAAAT | NM_003075.5 | Mission shRNA TRCN0000329883 |
| | shSMARCC2 #5 | GCCTGTCTCGACCTAACATTT | | Mission shRNA TRCN0000329885 |
| BRG1 | shBRG1 | TCAAACAGTACCAGATCAAAG | NM_003072 | Mission shRNA TRCN0000218544 |
| Non-Silencing Control | | CAACAAGATGAAGAGCACCAA | No target | Sigma-Aldrich SHC002 |
| ***siRNA*** | | | | |
| ARID1A | siARID1A | GCCTGATCTATCTGGTTCAAT | NM_006015.6 | Sigma-Aldrich custom, sequence from [21] |
| ARID1B | siARID1B | ACCATGAAGACTTGAACTTAA | NM_017519.3 | Sigma-Aldrich custom, sequence from [22] |

**Table S10**: Antibodies used for Western blot analysis.

| *Target protein* | *Origin* | *Dilution* | *Reference* | *Company* |
|---|---|---|---|---|
| ***Primary antibodies*** | | | | |
| SMARCA4/BRG1 | Mouse monoclonal | 1:1000 in 5% BSA | sc-17796 | Santa Cruz Biotechnology |
| SMARCB1/hSNF5 | Mouse monoclonal | 1:1000 in 5% milk | sc-166165 | Santa Cruz Biotechnology |
| ARID1A | Mouse monoclonal | 1:1000 in 5% milk | sc-32761 | Santa Cruz Biotechnology |
| ARID1B | Mouse monoclonal | 1:500 in 5% milk | ab57461 | Abcam |
| DPF2 | Rabbit monoclonal | 1:1000 in 5% milk | ab134942 | Abcam |
| BRD7 | Mouse monoclonal | 1:1000 in 5% milk | sc-376180 | Santa Cruz Biotechnology |
| BRD9 | Rabbit polyclonal | 1:500 in 5% milk | A303-781A-T | Bethyl Laboratories |
| SMARCC1/BAF155 | Mouse monoclonal | 1:1000 in 5% milk | sc-32763 | Santa Cruz Biotechnology |
| SMARCC2/BAF170 | Rabbit monoclonal | 1:1000 in 5% milk | #12760 | Cell Signaling Technology |
| PBRM1 | Mouse monoclonal | 1:1000 in 5% milk | AMAb90690 | Sigma-Aldrich |
| BRM | Rabbit polyclonal | 1:500 in 5% milk | ab15597 | Abcam |
| HDAC1 | Mouse monoclonal | 1:1000 in 5% milk | #5356 | Cell Signaling Technology |
| Cyclin D1/CCND1 | Rabbit monoclonal | 1:10000 in 5% BSA | ab134175 | Abcam |
| Phosphorylated-Rb | Rabbit monoclonal | 1:1000 in 5% BSA | #8516 | Cell Signaling Technology |
| p27 | Rabbit monoclonal | 1:2000 in 5% BSA | #3686 | Cell Signaling Technology |
| CASP3 (Full length) | Rabbit polyclonal | 1:2000 in 5% BSA | #9662 | Cell Signaling Technology |
| CASP3 Cleaved | Rabbit monoclonal | 1:500 in 5% BSA | #9664 | Cell Signaling Technology |
| PARP | Rabbit polyclonal | 1:2500 in 5% BSA | #9542 | Cell Signaling Technology |
| LOXL2 | Rabbit polyclonal | 1:1000 in 5% milk | NBP1-32954 | Novus Biologicals |
| NOTCH2 intracellular domain | Rabbit polyclonal | 1:500 in 5% milk | ab8927 | Abcam |
| Integrin β3/ITGB3 | Rabbit monoclonal | 1:1000 in 5% BSA | #13166 | Cell Signaling Technology |
| Integrin α9/ITGA9 | Mouse monoclonal | 1:1000 in 5% BSA | H00003680-M01 | Abnova |
| Phospho-FAK | Mouse monoclonal | 1:200 in 5% BSA | sc-81493 | Santa Cruz Biotechnology |
| FAK | Rabbit polyclonal | 1:1000 in 5% BSA | #3285 | Cell Signaling Technology |
| Phospho-SRC | Rabbit monoclonal | 1:1000 in 5% BSA | #6943 | Cell Signaling Technology |
| SRC | Rabbit polyclonal | 1:1000 in 5% BSA | #2108 | Cell Signaling Technology |
| ILK | Rabbit monoclonal | 1:1000 in 5% milk | ab52480 | Abcam |
| Actin | Mouse monoclonal-HRP | 1:20000 in 5% BSA | sc-47778 HRP | Santa Cruz |
| ***Secondary antibodies*** | | | | |
| Rabbit IgG | Goat polyclonal-HRP | 1:10000 in 5% BSA or milk | A0545 | Sigma-Aldrich |
| Mouse IgG | Rabbit polyclonal-HRP | 1:10000 in 5% BSA or milk | A9044 | Sigma-Aldrich |

Headquarters: Santa Cruz Biotechnology, Dallas, TX, USA; Abcam, Cambridge, UK; Bethyl Laboratories, Montgomery, TX, USA; Cell Signaling Technology, Danvers, MA, USA; Sigma-Aldrich, St. Louis, MO, USA; Novus Biologicals, Centennial, CO, USA; Abnova, Taipei, Taiwan.
